# Motor learning engages intrinsic plasticity in the human cortex

**DOI:** 10.64898/2026.11.19.767269

**Authors:** Benjamin Clennell, Binyu Luo, Kei Tung Lam, Yu Hu, Angel V Peterchev, Ricci Hannah

## Abstract

Learning is commonly explained by changes in synaptic strength. In animals, however, learning also engages intrinsic plasticity — changes in how readily neurons convert synaptic input into firing. Its contribution to human learning has been unclear, because conventional non-invasive measures of cortical excitability primarily reflect synaptic strength. Here we used controllable-pulse transcranial magnetic stimulation to estimate two biophysical indices of intrinsic excitability in human motor cortex, before and after motor training. Training produced signatures of increased intrinsic excitability, selectively in the cortical representation of the trained muscle, but not an untrained muscle. Individuals showing larger intrinsic excitability changes learned faster, independently of changes in motor-evoked potential amplitude, an index of synaptic excitability that increased following training but did not predict learning rate. An axon model reproduced the measured changes through small shifts in membrane excitability. These findings provide evidence that intrinsic plasticity is expressed in the intact human brain, and that it represents a behaviourally relevant component of learning-related cortical adaptation.

## Introduction

Learning and memory underpin our ability to acquire and retain new knowledge and skills. Motor skill learning is fundamental to adaptive behaviour and to recovering function after neurological injury, yet the neuronal adaptations that support it, and why some individuals learn faster than others, remain incompletely understood. Here we show that an often overlooked form of neuronal adaptation, regulation of intrinsic excitability, is engaged during human motor learning and predicts individual differences in learning rate.

The dominant framework for explaining learning-related behavioural change is synaptic, with extensive evidence that experience modifies synaptic strength and structure through mechanisms such as long-term potentiation, long-term depression, and dendritic spine remodelling^1–5^. Experience-dependent plasticity, however, is not restricted to the synapse. Neurons also adjust their intrinsic excitability, how readily they convert synaptic input into spiking output, through changes in the function and distribution of voltage-gated ion channels^6–8^. A growing body of theoretical and empirical work has argued that this form of adaptation, termed intrinsic plasticity, has been systematically overlooked, and that a complete account of learning requires moving beyond a synapse-centric view^9–11^.

Despite this, two decades of research into human motor learning has relied extensively on transcranial magnetic stimulation (TMS) measures that predominantly index synaptic plasticity. When applied over motor cortex, TMS elicits motor-evoked potentials (MEPs) in contralateral muscles, providing a non-invasive readout of corticospinal excitability. Many studies of human motor learning have reported increases in MEP amplitude after training^12–20^, which are typically thought to reflect synaptic facilitation within motor cortical circuits^21,22^. These MEP changes, however, reveal little about the intrinsic membrane excitability of the neurons being activated. Consequently, whether human motor learning engages intrinsic plasticity remains largely unknown.

Evidence from animal studies suggests that this may be an important omission. Motor learning-related intrinsic plasticity has been documented in motor cortex and cerebellar neurons^23–29^. In rodent primary motor cortex, motor training induces changes in the intrinsic excitability of layer II/III and V pyramidal neurons that accompany synaptic plasticity and track motor performance^25–27^. These findings raise the possibility that a behaviourally important component of human motor learning has remained hidden from conventional neurophysiological measurements.

Controllable-pulse TMS (cTMS) provides a way to probe intrinsic excitability non-invasively by varying pulse duration to estimate the strength–duration time constant (SDTC) and rheobase — biophysical parameters that are sensitive to axonal membrane excitability and voltage-gated ion channel function^30–34^. Unlike conventional MEP amplitude measures, these parameters are derived from how motor threshold varies with pulse duration. These measures primarily reflect intrinsic rather than synaptic excitability, providing a non-invasive means of testing whether intrinsic plasticity occurs in the human cortex.

Here, we tested whether motor learning engages intrinsic plasticity in the human motor cortex, and whether its magnitude relates to individual differences in learning. Healthy adults were trained on a rapid thumb abduction task. Before and after training, we used cTMS to estimate SDTC and rheobase in the cortical representations of the trained thumb muscle and an untrained hand muscle, alongside MEP amplitude as a conventional index of synaptic facilitation. We reasoned that if intrinsic plasticity contributes to motor learning, training should produce muscle-specific changes in SDTC and rheobase and the magnitude of these changes should covary with learning independently of changes in MEP amplitude.

We show that motor training induces changes consistent with increased intrinsic excitability in the trained cortical representation. These changes track individual differences in learning rate independently of conventional MEP facilitation, which itself was unrelated to learning. This identifies intrinsic plasticity as a behaviourally relevant component of human motor skill acquisition, extending a mechanism previously characterised only in animal models to the human brain.

## Results

### Behavioural performance

Participants performed ballistic thumb abduction movements, with the goal of maximising peak acceleration along a single, outward direction (Figure 1a,b). Two participants whose mean peak acceleration never exceeded their own baseline, in any practice block or at 1-h retention, were excluded from all analyses, as this indicated that they had not engaged with the task (Methods). This left N = 35 for neurophysiological analyses and N = 34 for behavioural and brain–behaviour analyses (one participant’s behavioural data were lost).

**Figure 1.**
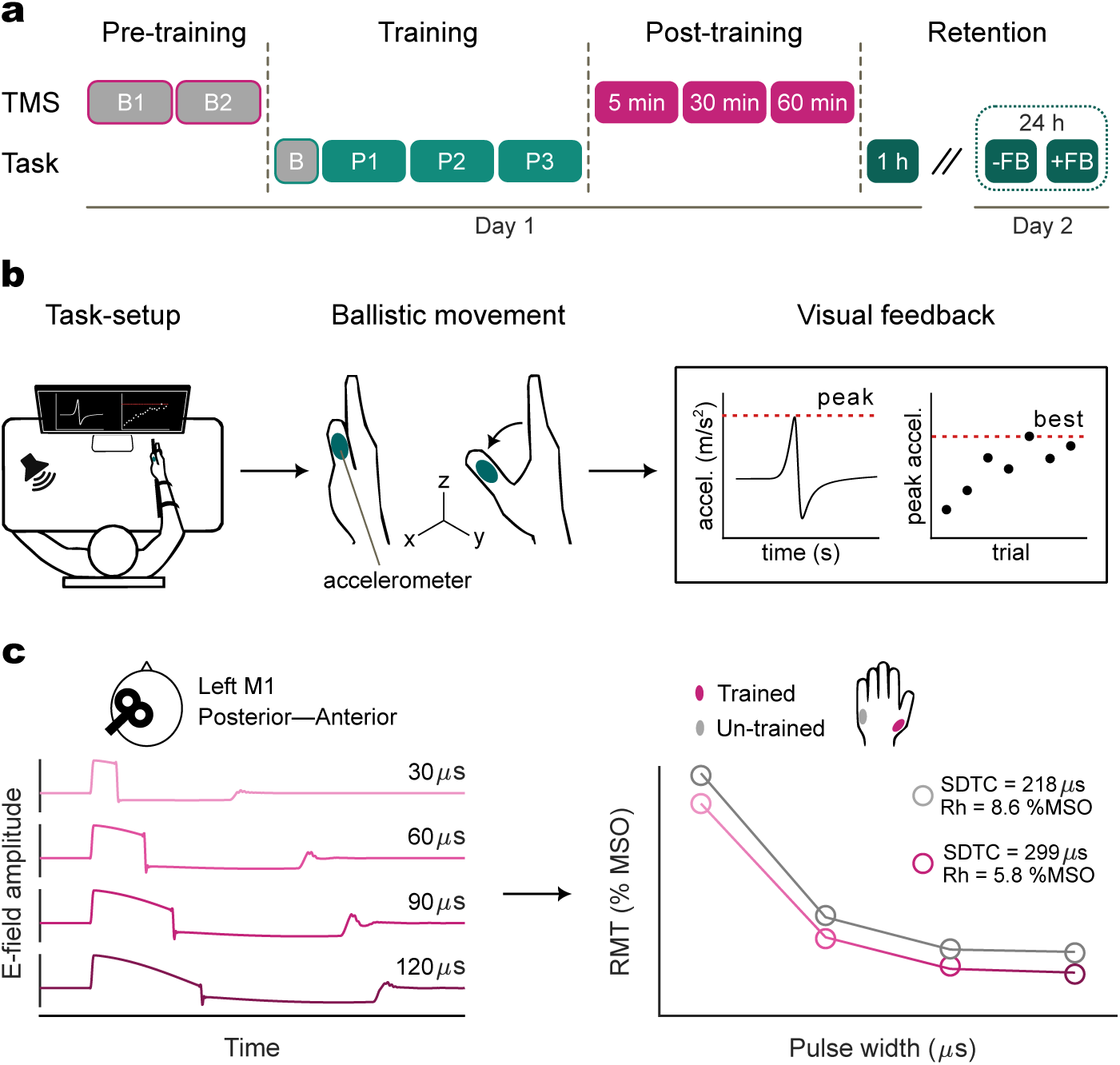
Experimental design and controllable-pulse transcranial magnetic stimulation (cTMS) methodology. (a) Experimental timeline. After two baseline cTMS assessments (B1, B2), participants completed the motor task across a baseline block (B; 20 trials) and three training blocks (P1–P3; 3 *×* 100 trials, separated by 5-min rests). cTMS assessments were repeated at 5, 30 and 60 min post-training. Behavioural retention was tested 1 h after the end of training (20 trials). A subset of participants (*n* = 17), completed two additional retention blocks 24 h later. In the first block, participants did not receive trial-by-trial feedback on performance (-FB). In the second block they did receive trial-by-trial feedback on performance (+FB). (b) Behavioural task. Participants are seated with their forearm fixed in a custom built rig to restrict movement of the hand and arm, leaving the thumb free to move. Movements are cued by a 1000 Hz tone paced at 0.33 Hz. Participants performed a ballistic thumb abduction with an accelerometer attached to the thumb, aiming to maximise peak acceleration along the abduction (*x*) axis. Participants received visual feedback of real-time *x*-axis acceleration on each trial (peak marked for illustration here only), alongside trial-by-trial tracking of peak acceleration, which was used to guide performance and reinforce learning across trials. (c) cTMS protocol. Stimulation was delivered over the left primary motor cortex (M1) in a posterior–anterior induced-current orientation using controllable-pulse TMS. Electric-field waveforms are shown for the four pulse widths used in the experiment (30, 60, 90 and 120 *µ*s). Resting motor threshold (RMT; % maximum stimulator output), was estimated by constructing input–output curves, defined as the intensity required to elicit a *∼*200 *µ*V MEP, and was measured across pulse widths for the trained APB and untrained ADM muscles. Fitting the strength–duration relationship yielded rheobase (Rh) and strength–duration time constant (SDTC). Strength–duration curves and the accompanying SDTC and Rh values are illustrative, from a single representative participant at baseline: open circles are measured RMT, solid lines are model fits.

Motor performance improved substantially over training, with mean peak acceleration increasing by 90% from baseline (Figure 2a; repeated-measures ANOVA: *F* (15, 495) = 13.83*, P <* 0.001*, η*^2^ = 0.30). The amount of learning varied across individuals (range: −23% to 452%), providing a basis for testing whether neurophysiological changes explained behavioural variability.

**Figure 2.**
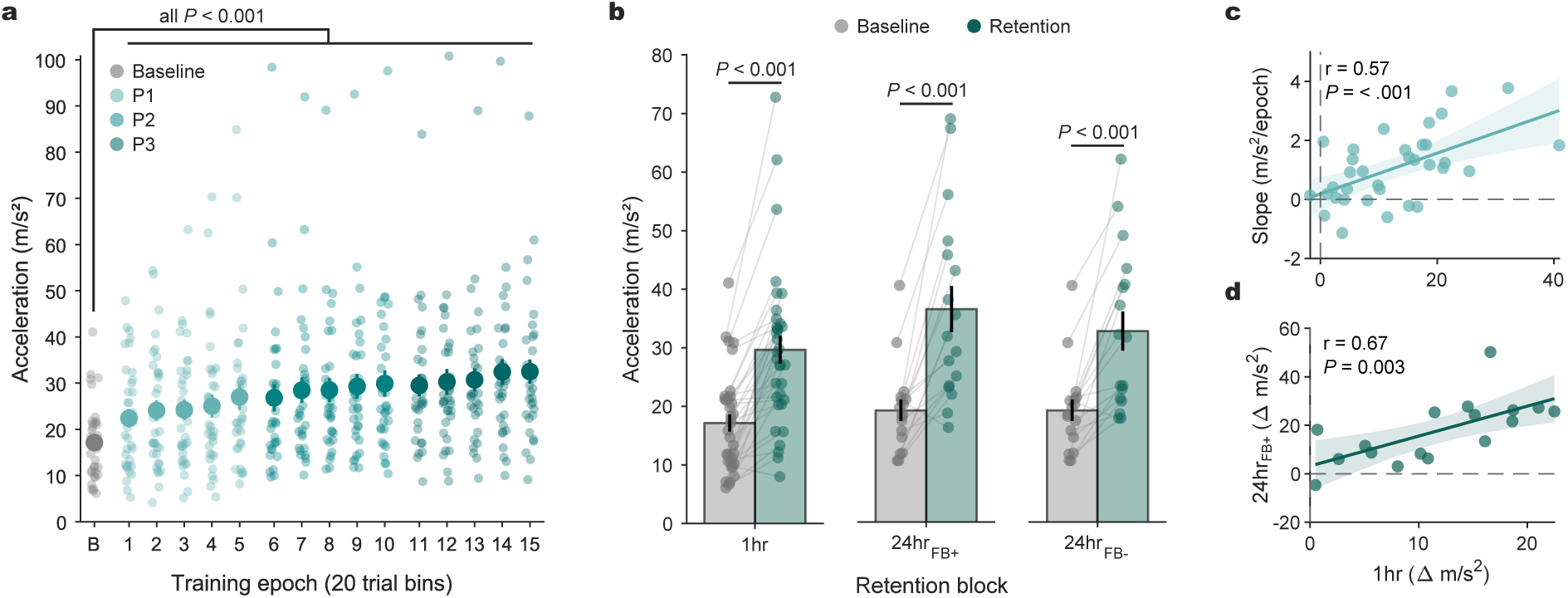
Ballistic thumb abduction training produces robust and durable improvements in task performance. (a) Group-mean peak thumb abduction acceleration (m/s^2^) across the baseline block and three successive practice blocks (100 trials each, binned into epochs of 20 trials). Peak acceleration increased across training, and all practice epochs exceeded baseline (all *P* < 0.001; paired t-test, Holm-corrected). Solid circles denote group mean; dots denote individual participants; error bars denote SEM; *N* = 34. (b) Peak acceleration remained above baseline during the 1-hour retention block (*N* = 34), and in the subsample (*N* = 17) tested at 24-hours without and with trial-by-trial visual feedback (all *P* < 0.001; paired t-test, Holm-corrected). (c) The relationship between mean within-block training slope and 1-hour retention (*r* = 0.57, *P* < 0.001). (d) The relationship between 1-hour retention and 24-hour retention (with feedback) in the subsample tested at both time points (*r* = 0.67, *P* = 0.003). In (a,b) data is mean, error is SEM, dots are individual subjects. In (c,d) dots represents individual subjects; the solid line denotes the linear fit and the shaded band the 95% confidence interval.

To quantify the rate of learning, we estimated the slope of peak acceleration across epochs within each practice block (P1–P3). The rate of learning did not differ across blocks (*F* (2, 66) = 2.16*, P* = 0.12), so their mean was used as the primary index of learning rate for subsequent analyses.

To assess short-term retention, participants completed a 20-trial block one hour after training. Performance remained above baseline (*P_adj_ <* 0.001; Figure 2b). In a subsample tested after 24 hours (*N* = 17), retention was first assessed without trial-by-trial performance feedback, to test whether performance gains reflected durable learning rather than online use of the feedback signal. Performance remained above baseline both without and with feedback (*P_adj_ <* 0.001; Figure 2b), indicating that the trained improvement was genuinely retained rather than dependent on concurrent feedback. Short- and long-term retention were closely related in this subset (Figure 2d), supporting the use of one-hour retention as the primary retention outcome for brain-behaviour analyses in the full sample. Learning rate and one-hour retention were moderately correlated, sharing roughly a third of their variance (*r* = 0.57*, R*^2^ = 0.32*, P <* 0.001; Figure 2c), so we treated these as related but separable behavioural indices of acquisition and retention in subsequent brain-behaviour analyses.

### Training-induced changes in intrinsic excitability

To assess whether motor learning was accompanied by changes in intrinsic excitability, we estimated SDTC and rheobase before training and at 5, 30 and 60 min afterwards (Figure 1a,c). Values were normalised to the group-mean pre-training value for each muscle.

SDTC and rheobase were stable across the two pre-training assessments in the trained muscle (SDTC: *P* = 0.30; rheobase: *P* = 0.43; paired *t*-tests). Differences were observed in the untrained muscle (SDTC: *P* = 0.018, *d* = *−*0.42; rheobase: *P* = 0.003, *d* = 0.54), so the average of both assessments was used as the pre-training reference. The following effects were preserved when only the immediate pre-training assessment was used (Supplementary Table 1–2).

Because our hypothesis concerned a change from pre- to post-training rather than its timing, the primary analysis compared pre-training values with the mean of the three post-training assessments, using a 2 *×* 2 (Muscle *×* Time) repeated-measures ANOVA. Training altered SDTC differently in the two muscles (Muscle *×* Time: *F* (1, 34) = 7.06, *P* = 0.012, *η*^2^_*p*_ = 0.17). SDTC increased in the trained muscle (+8.4%, *d* = 0.42, *P* = 0.017) but not in the untrained muscle (*−*1.2%, *d* = *−*0.06, *P* = 0.71; Figure 3a). Rheobase showed a complementary pattern (*F* (1, 34) = 6.51, *P* = 0.015, *η*^2^_*p*_ = 0.16), decreasing in the trained muscle (*−*6.7%, *d* = *−*0.41, *P* = 0.022) but not in the untrained muscle (*−*1.3%, *d* = *−*0.09, *P* = 0.59; Figure 3c). These effects were robust to outlier exclusion (Supplementary Table 1–2) and did not depend on the MEP criterion used to derive threshold (Supplementary Fig. S1).

**Figure 3.**
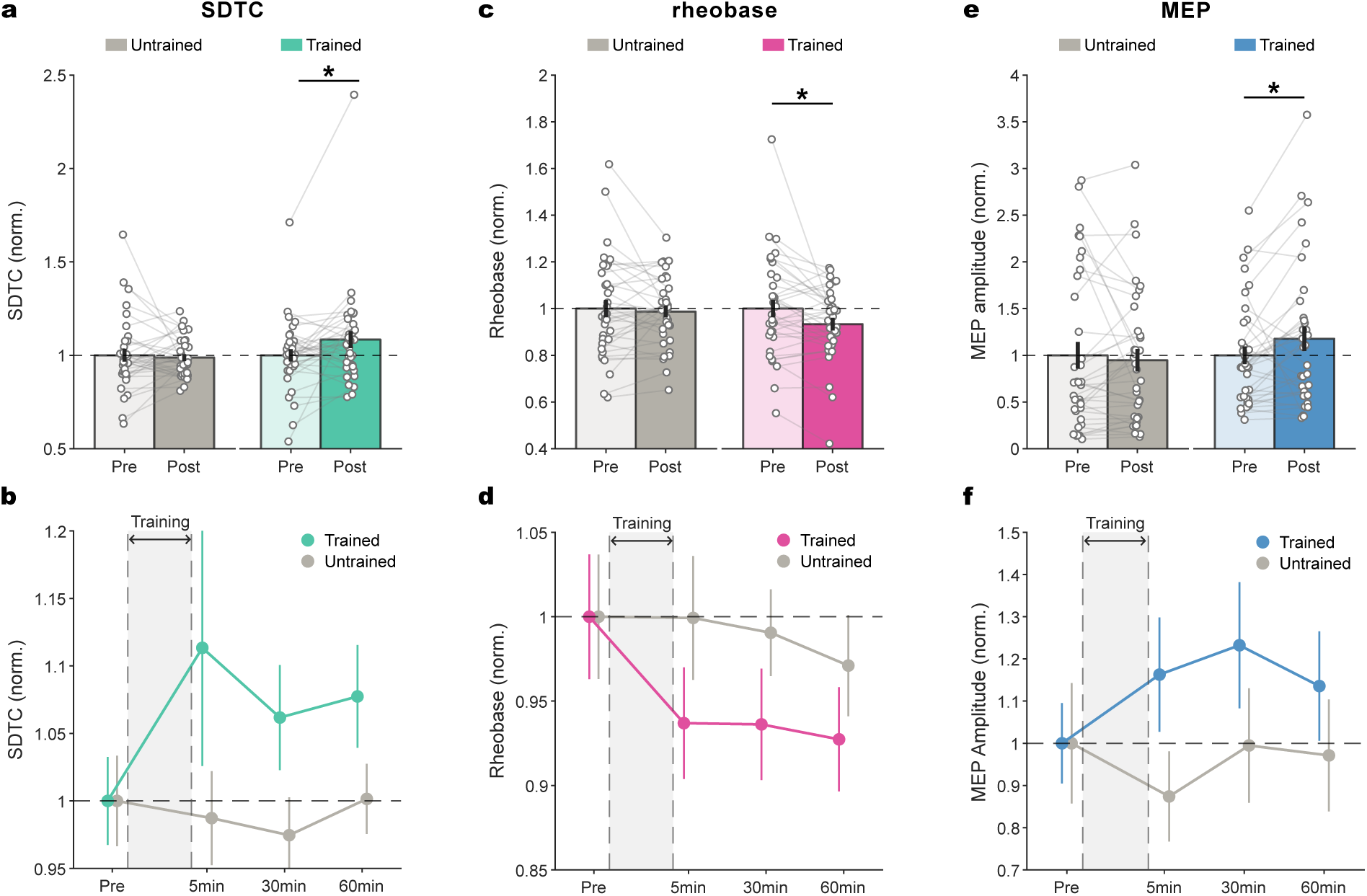
Motor learning induces muscle-specific changes in intrinsic excitability and MEP amplitude. Strength–duration time constant (SDTC; a,b), rheobase (c,d) and MEP amplitude (e,f) in the trained (APB) and untrained (ADM) muscles, normalised to the group-mean pre-training value for each muscle. (a) SDTC before and after training. Post-training values are the mean of the 5, 30 and 60 min assessments. SDTC increased in the trained but not the untrained muscle. (b) Time course of SDTC across the pre-training and three post-training assessments. (c) Rheobase before and after training. Rheobase decreased in the trained but not the untrained muscle. (d) Time course of rheobase. (e) MEP amplitude, averaged across pulse widths (30, 60, 90 and 120 *µ*s), before and after training. MEP amplitude increased in the trained but not the untrained muscle. (f) Time course of MEP amplitude, averaged across pulse widths. In (a,c,e), dots denote individual participants; \**P <* 0.05, paired *t*-test of pre-versus post-training values. Data are group means *±* SEM. *N* = 35.

We next examined the time course of these changes over the hour after training (Figure 3b,d) using a 2 *×* 4 ANOVA including all four time points. This showed a main effect of Muscle (SDTC: *F* (1, 34) = 4.37, *P* = 0.044, *η*^2^_*p*_ = 0.11; rheobase: *F* (1, 34) = 6.95, *P* = 0.013, *η*^2^_*p*_ = 0.17), but no main effect of Time (SDTC: *F* (3, 102) = 0.63, *P*_GG_ = 0.56; rheobase: *F* (3, 102) = 1.26, *P*_GG_ = 0.29) and no Muscle *×* Time interaction (SDTC: *F* (3, 102) = 1.38, *P*_GG_ = 0.26; rheobase: *F* (3, 102) = 1.39, *P*_GG_ = 0.25; Supplementary Table 1–2). Thus, there was no clear evidence that the time course of intrinsic excitability changes differed between muscles.

As a complementary analysis, we asked whether training altered the relationship between SDTC and rheobase, the two fitted parameters that jointly describe the strength–duration function. Training changed this relationship in the trained muscle but not the untrained muscle (Extended Data Fig. 1), providing converging evidence for a learning-related change in intrinsic excitability.

Because SDTC and rheobase are jointly estimated from motor thresholds across pulse widths, we next examined whether the training effect was also apparent when considering the thresholds themselves. We observed no significant change in motor threshold at any pulse width in either muscle (Extended Data Fig. 2). Forward modelling using each participant’s fitted SDTC and rheobase values predicted only sub-percent threshold changes. Thus, training altered the overall strength– duration profile, but this change would have been largely missed by a motor threshold measured at a single pulse width.

Together, these results show that motor learning produced complementary changes in SDTC and rheobase in the trained muscle, and that the relationship between these parameters also changed after training. Together, this provides evidence for a learning-related change in the intrinsic excitability of neurons in the cortical representation of the trained muscle.

### Training-induced changes in MEP amplitude

Alongside these intrinsic excitability measures, we examined MEP amplitude as a conventional index of training-related corticospinal facilitation. MEP amplitude was averaged across suprathreshold intensities and pulse widths (Methods). It was normalised to the group-mean pre-training value for each muscle and analysed in the same way as SDTC and rheobase.

Training altered MEP amplitude differently in the two muscles (Muscle *×* Time: *F* (1, 34) = 11.40, *P* = 0.002, *η*^2^_*p*_ = 0.25). MEP amplitude increased in the trained muscle (+17.7%, *d* = 0.36, *P* = 0.041) but not in the untrained muscle (*−*5.3%, *d* = *−*0.11, *P* = 0.53; Figure 3e). A 2 *×* 4 ANOVA including all four time points also showed a Muscle *×* Time interaction (*F* (3, 102) = 5.98, *P*_GG_ = 0.001, *η*^2^_*p*_ = 0.15), with no main effect of Muscle (*F* (1, 34) = 1.17, *P* = 0.29) or Time (*F* (3, 102) = 0.94, *P*_GG_ = 0.41; Figure 3f; Supplementary Table 4–5).

This pattern did not depend on pulse width. In a Time *×* Muscle *×* Pulse width ANOVA, the Time *×* Muscle interaction remained significant (*F* (3, 102) = 6.00, *P*_GG_ = 0.001, *η*^2^_*p*_ = 0.15) and did not interact with pulse width (*F* (9, 306) = 0.50, *P*_GG_ = 0.81; Supplementary Table 4). We therefore used the pulse-width-averaged change in MEP amplitude, averaged across post-training assessments, as the index of MEP facilitation in subsequent brain–behaviour analyses. This index is derived from the same input–output curves used to estimate SDTC and rheobase.

### Intrinsic excitability changes track behavioural learning independently of synaptic facilitation

To test whether training-induced neurophysiological changes were behaviourally relevant, we examined the relationship between changes in SDTC, rheobase, and MEP amplitude and two behavioural indices: learning rate and short-term retention.

In the trained muscle, changes in both intrinsic excitability parameters correlated with the learning rate. Greater increases in SDTC were associated with faster learning (*ρ* = 0.440, *P*_FDR_ = 0.015; Figure 4a) and greater decreases in rheobase were likewise associated with faster learning (*ρ* = *−*0.452, *P*_FDR_ = 0.015; Figure 4b). Change in MEP amplitude was unrelated to learning rate (*ρ* = 0.11, *P* = 0.53; Figure 4c). In the untrained muscle, no neurophysiological measure predicted learning rate (all *|ρ| ≤* 0.11, all *P >* 0.55; Figure 4d-f).

**Figure 4.**
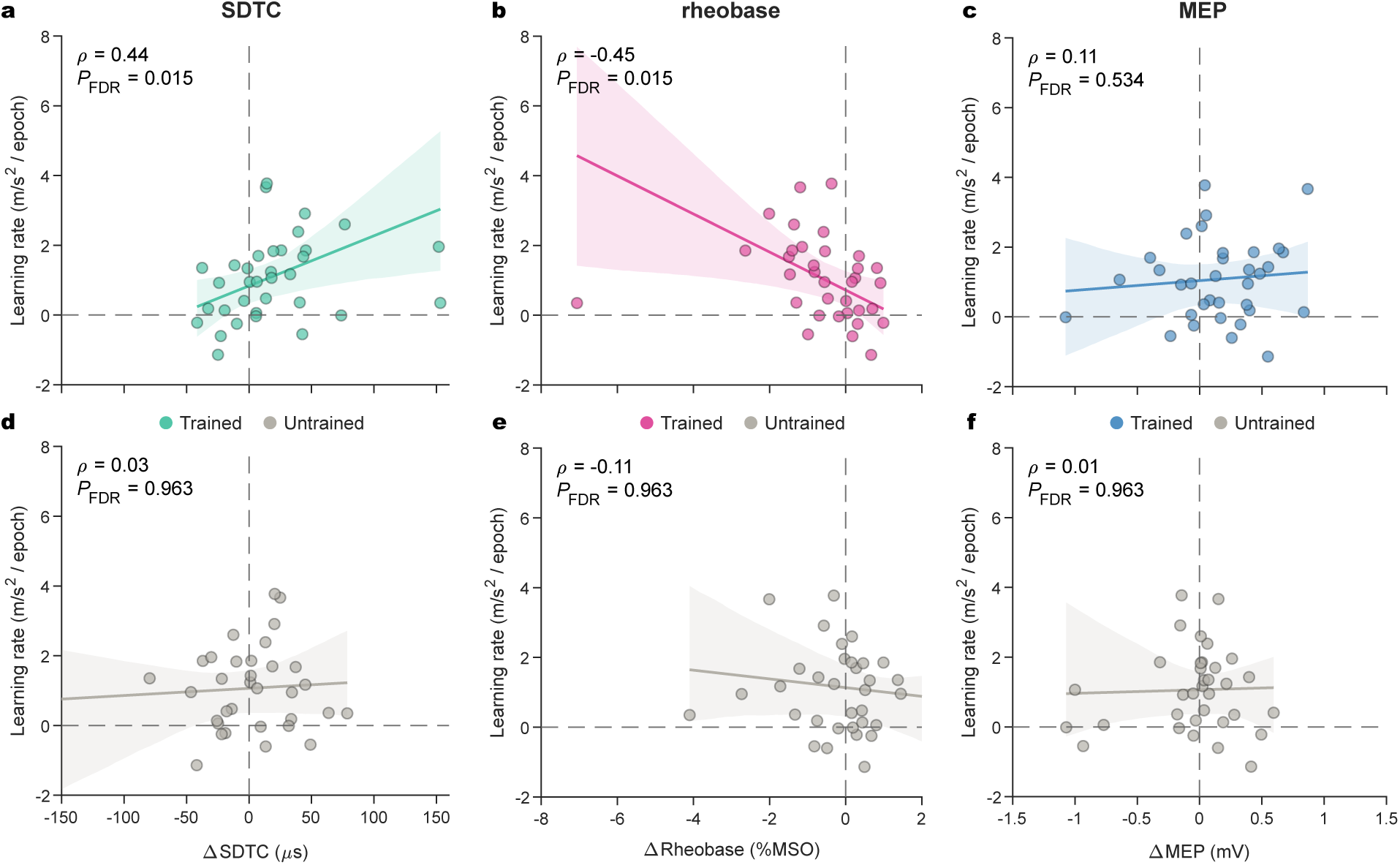
Muscle-specific intrinsic excitability changes predict the rate of learning during training. Spearman correlations between pre-to-post change in each neurophysiological measure, collapsed across the three post-training time points, and learning rate, quantified as the mean within-block training slope. Associations are shown for the trained muscle (top row) and untrained control muscle (bottom row). (a) Greater increases in trained muscle SDTC were associated with faster learning rates. (b) Greater decreases in trained muscle rheobase were associated with faster learning rates. (c) Change in trained muscle MEP amplitude (averaged across pulse widths) was unrelated to learning rate. (d, e, f) No neurophysiological measure in the untrained muscle was associated with learning rate. Each dot represents one participant; solid lines denote the Theil–Sen robust regression fit and shaded bands the 95% bootstrap confidence interval (10,000 resamples); *N* = 34.

To test whether the relationship between intrinsic changes and learning rate were independent of corticospinal facilitation, we computed partial correlations controlling for changes in MEP amplitude. Both intrinsic excitability associations remained significant after controlling for MEP change (SDTC: *ρ*_partial_ = 0.47*, P* = 0.006; Rheobase: *ρ*_partial_ = *−*0.44*, P* = 0.010; Extended Data Fig. 3a,d). In contrast, MEP change did not predict learning rate after controlling for either SDTC (*ρ*_partial_ = 0.21*, P* = 0.25; Extended Data Fig. 3b) or rheobase (*ρ*_partial_ = 0.04*, P* = 0.84; Extended Data Fig. 3e). Changes in SDTC and rheobase were also not significantly correlated with changes in MEP amplitude within the trained muscle (SDTC–MEP: *ρ* = *−*0.17*, P* = 0.35; rheobase– MEP: *ρ* = *−*0.17*, P* = 0.33; Extended Data Fig. 3c,f), supporting a dissociation between intrinsic excitability changes and conventional MEP facilitation. Sensitivity analyses showed consistent patterns across alternative analysis choices (Supplementary Table 6–8).

Associations with short-term retention were weaker than those observed for learning rate. In the trained muscle, SDTC and rheobase changes were in the same direction as the learning rate findings, but neither association was significant (SDTC: *ρ* = 0.21, *P* = 0.23; rheobase: *ρ* = *−*0.29, *P* = 0.099; Extended Data Fig. 4a,b). MEP amplitude change was unrelated to retention (*ρ* = *−*0.04*, P* = 0.84; Extended Data Fig. 4c). No neurophysiological measure from the untrained muscle was associated with retention (all *|ρ| ≤* 0.11, all *P >* 0.52; Extended Data Fig. 4d–f). In the subsample retested at 24 h (*N* = 17), no measure was associated with long-term retention in either muscle (all *P >* 0.16; Supplementary Table 9).

Overall, changes in intrinsic excitability — but not MEP facilitation — tracked individual differences in learning rate.

### An axon model supports a membrane-based interpretation of the strength–duration changes

The preceding results show that training increased SDTC and decreased rheobase in a muscle-specific manner, and that the magnitude of these changes scaled with learning. SDTC and rheobase are biophysical indices of axonal excitability, but the observed changes do not uniquely identify the underlying membrane mechanism. To provide an additional layer of mechanistic plausibility, we used a biophysically detailed model of a myelinated axon to test whether realistic changes in ion-channel conductance or resting membrane potential could reproduce the observed combination of increased SDTC and decreased rheobase. Because the fitted strength–duration profile could also change if training altered the population of TMS-sensitive axons contributing to the MEP, we additionally varied axon diameter as a first-order proxy for a change in recruitment geometry. The aim was not to identify a unique mechanism, but to test whether the experimental signature was compatible with plausible membrane-level changes and whether it could be reproduced by a simple shift in the morphology of the recruited axons.

We simulated a straight, myelinated axon using the MRG double-cable model in the small-calibre refit of Peña and colleagues^35^, which extends the model’s validity to fibres of approximately 1 *µ*m to 6 *µ*m and so to the calibre range reported for myelinated cortical axons^36,37^. Major intracortical axon collaterals in motor cortex, the elements most likely activated by threshold PA-TMS, are myelinated with axon diameters of 1–3 *µ*m^38^; we took a fibre whose internodal axon diameter lies at the centre of this range (2 *µ*m; 3.66 *µ*m fibre diameter) as the baseline. We stimulated the model with the measured cTMS waveforms and estimated SDTC and rheobase from the simulated thresholds using the same membrane model applied to the experimental data. The model’s baseline SDTC (284 *µ*s) fell within the range of experimentally observed baseline values (group mean 229 *µ*s, range 120–387 *µ*s). Although a peripheral fibre model, its nodal membrane description corresponds to that used as the nominal neuron model in TMS pulse-dynamics work, and threshold behaviour across pulse widths is relatively insensitive to the specific neuron model adopted^39,40^; we therefore treat it as a generic account of myelinated axon excitation.

We varied five parameters independently and examined how each altered SDTC and rheobase (Figure 5a–e). Because sensitivity differs markedly between parameters, each sweep was scaled to span changes in SDTC and rheobase of the order observed experimentally. Changes in persistent sodium conductance, resting membrane potential and slow potassium conductance reproduced the experimentally observed combination of SDTC and rheobase change most closely: when matched to the observed median rheobase decrease, each produced an SDTC increase of 4.2%, compared with 5.3% experimentally (Figure 5f). Transient sodium conductance produced a smaller SDTC increase of 3.5%, and axon diameter only 1.1%. A change in the calibre of the activated axons therefore accounts least well for the coupled change. Thus, plausible changes in membrane excitability readily generated the experimental pattern, whereas a simple shift in the geometry of the recruited axon did not.

**Figure 5.**
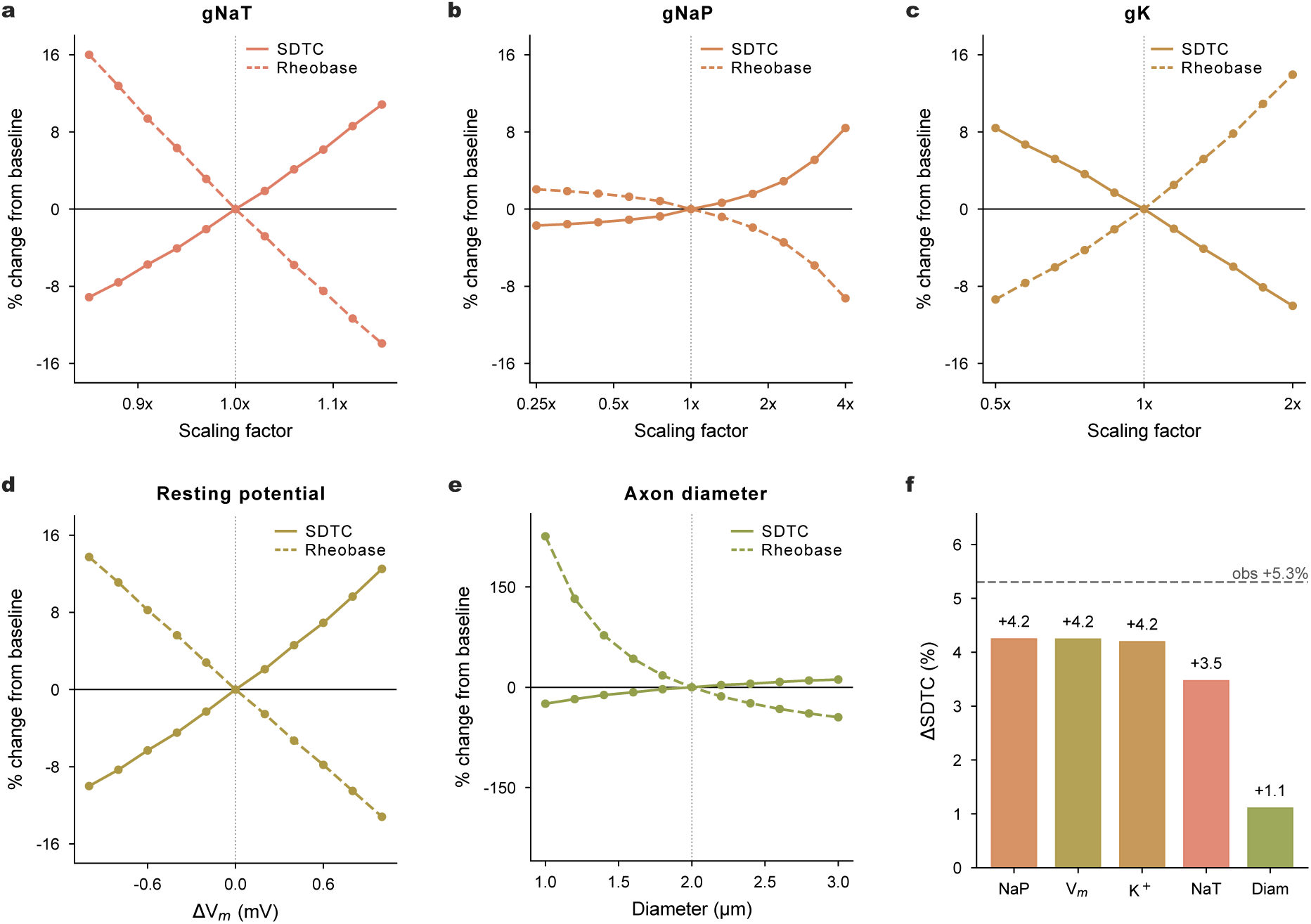
An axon model identifies candidate membrane mechanisms for the observed strength–duration changes. A straight myelinated model axon was stimulated with the measured cTMS waveforms at four pulse widths (30, 60, 90, 120 *µ*s). SDTC and rheobase were estimated from the simulated thresholds using the same strength–duration model applied to the experimental data. (a–e) Effect of varying five biophysical parameters independently on model SDTC (solid lines) and rheobase (dashed lines), expressed as percentage change from baseline: (a) transient sodium conductance (*g*NaT), (b) persistent sodium conductance (*g*NaP), (c) slow potassium conductance (*g*K), (d) resting membrane potential (*V*_m_) and (e) axon diameter. Dotted vertical lines mark baseline values. Conductance and resting-potential ranges were chosen to span the observed SDTC and rheobase changes. Axon diameter spanned the range reported for the myelinated intracortical axon collaterals likely activated by posterior–anterior TMS38. Panel (e) uses a wider y-axis range. (f) Change in SDTC at the parameter value that reproduced the observed median within-subject rheobase change (*−*4.9%). The dashed line marks the observed median within-subject SDTC change (+5.3%). Enhanced *g*NaP, reduced *g*K and depolarisation of *V*_m_ reproduced the observed SDTC increase most closely.

Because altering a nodal conductance also shifts the membrane’s resting potential, we measured the resting potential at the node for each matched condition. At their matched values, the three best-fitting parameters all depolarised the node by a similar amount (persistent sodium +0.42 mV, resting potential +0.37 mV, slow potassium +0.29 mV), whereas transient sodium conductance and axon diameter produced little or no depolarisation (+0.08 and +0.02 mV). These parameters therefore converge on a common effect on resting membrane excitability.

## Discussion

The present study provides evidence that motor learning engages intrinsic plasticity in the human motor cortex. Using controllable-pulse TMS before and after motor training, we found increases in SDTC and decreases in rheobase specifically in the trained motor representation. To our knowledge, this provides the first non-invasive demonstration of learning-related intrinsic plasticity in the human cortex, identifying a form of neuronal adaptation that has been central to animal accounts of learning but difficult to study directly in humans.

The learning-related increase in intrinsic excitability observed here is consistent with cellular evidence from animal models28,41–46. Learning has been associated with lower current thresholds for spike generation in cat motor cortex neurons23,24, while more recent work in identified rodent engram neurons has shown coordinated changes in rheobase and membrane dynamics47. Manipulating the excitability of these neurons altered the strength and precision of subsequent memories, providing causal evidence that intrinsic excitability can influence memory formation. Intrinsic plasticity is also temporally dynamic: in rat motor cortex, it can emerge within minutes of conditioning48 and evolves over days of training25,26. The within-session changes measured here may therefore represent an early phase of a longer plasticity trajectory.

A central finding was that the intrinsic excitability changes were behaviourally relevant. Individuals who showed larger increases in SDTC and larger decreases in rheobase in the trained motor representation learned faster, and these relationships remained after accounting for corticospinal facilitation. Animal work has led to the proposal that transient increases in intrinsic excitability create a hyperexcitable state that biases which neurons are recruited into a memory ensemble and primes them for subsequent consolidation9,10. Our findings are consistent with this account and extend it to human motor learning by showing that learning-related intrinsic excitability changes covary with individual differences in the rate of learning.

The behavioural relevance of the intrinsic excitability changes was dissociable from conventional MEP facilitation. Although motor training produced muscle-specific MEP facilitation alongside the SDTC and rheobase changes, MEP change did not correlate with learning rate, and the intrinsic excitability-learning relationships remained after controlling for MEP changes. MEP facilitation has been the dominant readout of learning-related M1 plasticity in humans, typically interpreted as reflecting LTP-like synaptic change12,13,49–51. Yet associations between training-related MEP changes and behavioural learning have been reported only occasionally12,14, with most studies finding no clear relationship13,15–20. Our findings therefore suggest that MEP amplitude alone provides an incomplete account of the cortical adaptations supporting skill acquisition. By characterising strength–duration properties, SDTC and rheobase captured a distinct, behaviourally relevant component of learning-related cortical plasticity.

The coordinated increase in SDTC and decrease in rheobase provides a biophysically coherent signature of altered membrane excitability. In our axon model, enhanced persistent sodium conductance, reduced slow potassium conductance and a small depolarisation of the resting potential each reproduced this signature to a similar degree. Notably, all three manipulations produced a comparable subthreshold depolarisation (0.3–0.4 mV) of the axon. These routes are therefore unlikely to be wholly independent. Persistent sodium current activates in the subthreshold range, so a depolarisation of this magnitude increases its availability52. These findings suggest that a small shift in resting membrane potential, potentially accompanied by increased persistent sodium current, could underlie the observed changes in SDTC and rheobase.

In peripheral axons, where strength–duration properties have been characterised most extensively, SDTC is largely attributed to persistent sodium current at the node52–56, and depolarising interventions such as ischaemia increase SDTC through this current57. Pharmacological evidence in the human motor cortex supports this: lacosamide, which promotes slow inactivation of voltage-gated sodium channels and reduces persistent sodium current58,59, produces the opposite pattern of SDTC and rheobase change in both cortical33 and peripheral motor axons60. Persistent sodium current is itself a target of neuronal plasticity: it has been implicated in associative memory and memory-relevant coding61–63, and activity-dependent plasticity can modify sodium-channel function and localisation64–66. Potassium conductance and resting membrane potential nonetheless remain plausible contributors, and are often implicated as intrinsic plasticity mediators in animals25–27,41,42,44,47. These accounts are complementary rather than competing, and neither the data nor the modelling uniquely identify the mechanism; resolving this will require pharmacological cTMS studies with agents having defined actions on sodium and potassium conductances.

An alternative interpretation is that training changed the population of TMS-sensitive neural elements contributing to the criterion MEP, rather than the intrinsic properties of the same population. For example, training-related synaptic strengthening could allow a previously weakly contributing population of TMS-sensitive inputs to contribute more strongly to the MEP and thereby alter the fitted strength–duration parameters. Two observations argue against this account. First, it predicts some dependence on MEP facilitation, yet MEP amplitude change was unrelated to learning rate and to the SDTC and rheobase changes, and controlling for MEP change did not attenuate the intrinsic excitability–behaviour relationships. Second, changing axon diameter in our model — a first-order proxy for a shift in recruitment geometry — predominantly altered rheobase and did not reproduce the coupled increase in SDTC and decrease in rheobase. Together these observations favour the interpretation that motor practice altered the intrinsic excitability of TMS-sensitive cortical elements.

Although demonstrated here in motor cortex, intrinsic plasticity is unlikely to be unique to motor learning. Learning-related intrinsic excitability changes are documented across the hippocampus41–43, amygdala45,67, prefrontal cortex47,68, sensory cortex69, and cerebellum28,70, and the voltage-gated ion channel mechanisms that support them are widely expressed throughout the brain. Intrinsic plasticity is therefore likely to operate wherever cortical circuits encode new information. The present findings therefore raise the possibility that comparable adaptations occur during other forms of human learning. Testing this possibility will require methods that can probe intrinsic excitability without relying on a peripheral motor response.

Intrinsic excitability may also be a key determinant of how cortical circuits respond to neuro-modulatory interventions such as repetitive TMS71,72, transcranial ultrasound stimulation73–75, and transcranial direct current stimulation76,77. These approaches are increasingly explored as adjuncts to motor training and neurorehabilitation, as well as for modifying cognition and behaviour more broadly. Measures that distinguish intrinsic from synaptic excitability could therefore help explain variability in response, identify when stimulation engages learning-relevant mechanisms and support development of more mechanistically informed interventions.

These findings provide non-invasive evidence that intrinsic plasticity is engaged during human motor learning. They identify changes in intrinsic excitability as a behaviourally relevant component of skill acquisition, dissociable from the MEP-based measures of facilitation that have dominated human motor plasticity research. These results bring a mechanism established in animal models into non-invasive human neuroscience, identifying intrinsic excitability as a measurable and behaviourally relevant component of skill acquisition.

## Methods

### Participants

Thirty-seven healthy young adults (23 female; mean age 23.1 years, SD = 5, range 18–36) completed the study. All participants provided written informed consent prior to testing, and all procedures were performed in accordance with the Declaration of Helsinki. The study was approved by the Research Ethics Committee at King’s College London (Project ID: HR-24/25-39903).

Because the study was designed to test for training-induced neurophysiological change, the analyses required that participants had engaged with the training task. Two participants whose mean peak acceleration never exceeded their own baseline, in any of the three practice blocks or at 1-h retention, were therefore excluded from all analyses; both remained 5–44% below baseline throughout. This criterion was deliberately conservative. It excluded only participants who showed no improvement at any point, and retained those who improved little or whose performance declined later in training: 7 of the remaining 34 participants had a negative mean learning-rate slope. One further participant’s behavioural data were lost due to a file save error. The analysed samples were therefore N = 35 for neurophysiological analyses (21 female, 14 male; mean age 23.3 years, SD = 5.1) and N = 34 for behavioural and brain–behaviour analyses, of whom 17 completed the 24-h retention session.

### Experimental design

Participants attended a single laboratory session in which they performed a ballistic thumb abduction task (300 training trials). Baseline cTMS measurements were acquired prior to training, and three post-training assessments were acquired at approximately 5, 30 and 60 min after the final training trial. A behavioural retention block was then performed at 1 h post-training. A subsample of participants (*n* = 17) returned 24 h later for an additional retention assessment.

### Ballistic thumb abduction task

#### Apparatus and setup

Participants were seated with their dominant arm slightly abducted, the elbow flexed to *∼*45*^◦^* (where 0*^◦^* is full extension) and the forearm semi-pronated. The wrist, hand and digits were secured in a custom-built rig that prevented movement at all joints except the thumb. Thumb acceleration was measured along the *x*, *y* and *z* axes using a tri-axial accelerometer (EVAL-ADXL326Z; Analog Devices Inc., USA) attached to the proximal phalanx of the thumb and aligned so that the abduction–adduction axis was equivalent to the *x*-axis of the accelerometer. The raw signal was digitised at 1000 Hz (NI USB-6002, National Instruments, USA), and stored for offline analysis. Task presentation and data acquisition were controlled using custom MATLAB R2024a software (MathWorks, USA).

#### Training protocol

The task was adapted from previous ballistic thumb abduction paradigms15,16,78,79. Participants were instructed to perform maximal-acceleration ballistic thumb abductions of the dominant hand, paced at 0.33 Hz by a brief auditory cue (1000 Hz tone). The session consisted of a baseline block of 20 trials followed by three training blocks of 100 trials each. Training blocks were separated by 5 min rest periods to minimise fatigue. During all blocks, participants received on-line visual feedback showing the real-time *x*-axis acceleration for each trial and trial-by-trial peak acceleration tracking with a persistent marker indicating the greatest acceleration achieved through-out training. The experimenter verbally encouraged participants to exceed their best performance throughout training.

#### Retention assessments

Retention was assessed at 1 h post-training using a single block of 20 trials, with feedback identical to that provided during training. The subsample of participants returning at 24 h (*n* = 17) completed two additional blocks of 20 trials: a first block without trial-by-trial feedback of peak acceleration, followed by a second block with feedback restored to match the training conditions.

### Electromyography

Surface electromyographic (EMG) activity was recorded from the abductor pollicis brevis (APB; Trained) and abductor digiti minimi (ADM; Untrained) of the dominant hand, with active electrodes placed over the muscle belly and reference electrodes over the proximal phalanges. A ground electrode was placed over the radial styloid process. Signals were amplified (*×*1000; Digitimer D360, Digitimer, UK), bandpass filtered (10–2000 Hz), passed through a 50 Hz noise eliminator (Hum Bug,

Digitimer, UK), and digitised at 5000 Hz (NI USB-6002, National Instruments, USA). Signals were acquired using custom MATLAB R2024a software.

### Transcranial magnetic stimulation

Single-pulse TMS was used to evoke motor-evoked potentials (MEPs) simultaneously from the APB (task-agonist) and ADM (task-uninvolved control) muscles of the dominant hand, delivered to the contralateral primary motor cortex (M1) before and at approximately 5, 30 and 60 min after training.

#### Stimulator and pulse waveforms

Stimulation was delivered using a controllable-pulse TMS device (cTMS3) connected to a 70 mm figure-of-eight coil (Elevate TMS, Rogue Research Inc., Montreal, Canada). The cTMS3 device generates near-rectangular induced electric-field waveforms with a large-amplitude initial phase followed by a low-amplitude, longer-duration second phase (Fig. 1c). The ratio of these amplitudes (M-ratio) was set to 0.15 for all pulses, so that the induced field was strongly unidirectional. Four pulse widths were used, referring to the duration of the initial positive phase: 30, 60, 90 and 120 *µ*s. The maximum pulse amplitude at each pulse width was limited by the device to 100%, 72%, 54% and 49% of maximum stimulator output (MSO) respectively.

#### Coil positioning

The coil was positioned over the M1 contralateral to the dominant hand and oriented *∼*45*^◦^* postero-laterally so as to induce posterior–anterior initial currents across the anterior bank of the central sulcus. The motor hotspot was defined as the scalp location at which slightly suprathreshold 120 *µ*s pulses produced the largest and most consistent MEP in APB, and was marked on a snug-fitting cap worn by the participant.

#### Threshold estimation and input–output curves

At each pulse width, a preliminary estimate of resting motor threshold (RMT) for the APB muscle was obtained using the stochastic approximation (SA) method80,81 with a response criterion of *≥*50 *µ*V peak-to-peak. These preliminary RMTs were used to set the stimulus intensities for input–output (I–O) curves, which characterise MEP amplitude as a function of stimulus intensity. For each pulse width, 8 pulses were delivered at each of five intensities (95%, 100%, 105%, 110% and 120% of preliminary RMT), with an additional 8 pulses at the maximum stimulator output for that pulse width. The intensity range was selected to encompass the 200 *µ*V MEP amplitude used as the criterion for estimating motor threshold (see *Input–output curve fitting*). The order of pulse width and intensity was randomised, and the inter-stimulus interval varied between 3 and 5 seconds.

Thresholds were derived from the I–O curves rather than measured directly, so that all outcome measures came from the same set of pulses. Threshold-tracking methods converge on the threshold of a single target muscle. Deriving thresholds from I–O curves instead allowed thresholds for both APB and ADM, and therefore SDTC and rheobase, to be estimated together with suprathreshold MEP amplitude from the same pulses, delivered at the same time.

### Data and statistical analyses

All data processing and statistical analyses were performed in MATLAB R2024a (MathWorks, USA). Standardised effect sizes are reported alongside all inferential tests: Cohen’s *d_z_* for paired comparisons and partial eta-squared (*η*^2^_*p*_) for omnibus ANOVA effects. The threshold for statistical significance was set at *α* = 0.05. Multiple-comparison correction was applied using the Holm– Bonferroni procedure for pairwise contrasts and the Benjamini–Hochberg false discovery rate (FDR) procedure for families of brain–behaviour correlations. Correction families were defined within each muscle throughout, as the trained and untrained muscles address distinct hypotheses: whether training altered a given measure, and whether any such change was anatomically specific.

#### Behavioural data processing

Trials containing equipment artefacts (e.g. voltage spikes) were identified by visual inspection and excluded. For each trial, acceleration along the primary movement axis was baseline-corrected by subtracting the mean acceleration over the initial 100 ms of the trial, removing any static offset due to sensor orientation relative to gravity. Peak acceleration was then identified in a two-step procedure: the global minimum acceleration (corresponding to movement deceleration) was first located, and the maximum positive acceleration was then identified within the 700 ms window preceding this negative peak. Trial-level peak accelerations were averaged into non-overlapping epochs of 20 trials for statistical analysis.

#### Training performance

Training-related changes in peak acceleration were assessed using a oneway repeated-measures ANOVA on epoch-level mean peak acceleration, with epoch as the within- subjects factor (spanning baseline and all training epochs). Where the omnibus test was significant, post-hoc paired t-tests compared each training epoch against baseline with Holm–Bonferroni correction. To characterise individual differences in the learning rate during training, ordinary least-squares linear slopes were fitted to the epoch-level mean peak acceleration within each of the three practice blocks. For the second and third practice blocks (P2, P3), slopes were fitted to the five within-block epochs only. For the first practice block (P1), the baseline epoch was included as epoch 0 and the five training epochs as epochs 1–5, so that the P1 slope captured the initial rate of change from pre-training performance. Because within-block slopes did not differ statistically across practice blocks (see Results), the mean of the three within-block slopes was used as the primary index of learning rate for brain–behaviour analyses, integrating learning across the full training period.

#### Retention

Retention was assessed using paired *t*-tests comparing baseline peak acceleration to each of three retention timepoints (1 h post-training, 24 h without feedback, 24 h with feedback), with Holm–Bonferroni correction applied across these three tests. The relationship between 1 h and 24 h retention was quantified using Pearson’s *r* with an ordinary least-squares fit. Cross-measure associations between learning rate and retention were also quantified using Pearson’s *r* to assess the degree of shared variance between acquisition rate and retention outcomes.

#### EMG preprocessing

Residual mains noise at 50 Hz was removed from each EMG trial using a least-squares fitting approach. The 500 ms signal segment immediately preceding the TMS pulse was used to estimate the amplitude and phase of a 50 Hz sinusoid, which was then subtracted from the entire trial. A baseline correction was subsequently applied by subtracting the mean signal amplitude of a 50 ms pre-stimulus window. Trials were excluded from all subsequent analyses if the peak-to-peak EMG amplitude within the pre-stimulus window exceeded 30 *µ*V.

#### Input–output curve fitting

Peak-to-peak MEP amplitudes were computed as the maximum peak-to-peak deflection in a 35 ms window (15–50 ms) following the TMS pulse. Amplitudes were log-transformed and a four-parameter Boltzmann sigmoid was fitted to the log-transformed MEP amplitude versus stimulus intensity (SI, % MSO) using non-linear least squares:

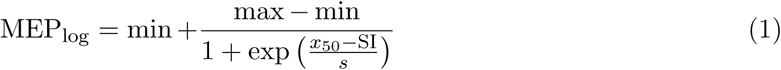

where MEP_log_ is the log-transformed MEP amplitude, *x*_50_ is the stimulus intensity producing a half-maximal response, and *s* governs the transition slope. The lower (min) and upper (max) plateaus were fixed across pulse widths within each session: min was set as the 5th percentile of the pre-stimulus EMG amplitude in a 50 ms pre-TMS window, and max as the 95th percentile of the observed peak-to-peak MEP amplitudes, both computed across all pulse widths and stimulus intensities within a given measurement block for a given participant. Curves were fitted for each combination of participant, block and pulse width. An I–O-derived resting motor threshold (RMT_I–O_) was defined as the stimulus intensity at which the fitted curve crossed log10(200), corresponding to a 200 *µ*V MEP. To confirm that the primary SDTC and rheobase changes were not dependent on the chosen amplitude criterion, thresholds were re-derived with 50 *µ*V, 100 *µ*V and 300 *µ*V criteria.

#### SDTC and rheobase estimation

The cortical strength–duration time constant (*τ_m_*, SDTC) and rheobase (*V*_th_*_,∞_*) were estimated from the set of RMT_I–O_ values across the four pulse widths (30, 60, 90, 120 *µ*s) using the leaky-integrator membrane model30,31,33,82, which incorporates the measured induced E-field waveform for each pulse configuration. The predicted threshold for a given pulse width *t_*p*_* is:

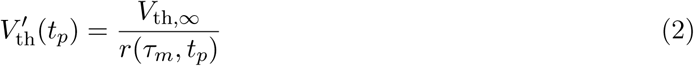

where *r*(*τ_m_, t_*p*_*) is the peak output of the leaky-integrator in response to the measured waveform. SDTC and rheobase were estimated per participant and block by minimising the sum of squared relative residuals between predicted and measured thresholds:

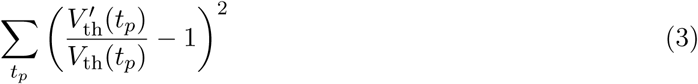

#### Neurophysiology data normalisation and outlier screening

A common normalisation scheme was applied to all neurophysiological measures (SDTC, rheobase, MEP amplitude) for analyses of training-induced change. For each measure and muscle, values were divided by the group grand mean at baseline (average of the two pre-training blocks), so that baseline mean equalled 1 and post-training values expressed proportional change. This grand-mean normalisation preserves between-subject variance while removing absolute scale differences between muscles and, for MEPs, between pulse widths. Participants with extreme change scores were identified using a median absolute deviation (MAD) criterion: for each muscle and measure, the change score (mean of the three post-training blocks minus baseline) was screened and participants exceeding 3.5*×* the scaled MAD (scaling constant 1.4826, for consistency with a normal distribution) were flagged. All primary analyses were repeated with flagged participants excluded, with the reduced sample re-normalised to its own grand-mean baseline so that sensitivity analyses were not biased by excluded participants’ baseline values.

#### Analysis of SDTC and rheobase

Our hypothesis was that training would change SDTC and rheobase from pre- to post-training in the trained but not the untrained muscle. This was tested with a 2 *×* 2 repeated-measures ANOVA with factors Time (pre, post) and Muscle (trained, untrained), in which post-training values were the mean of the 5, 30 and 60 min assessments. The Time *×* Muscle interaction was followed by paired *t*-tests of pre-against post-training values in each muscle. To characterise the time course, we ran a 2 *×* 4 ANOVA including all four time points (pre, 5, 30, 60 min). ANOVAs were implemented with MATLAB’s fitrm/ranova, with Greenhouse–Geisser correction where sphericity was violated. All tests were repeated using only the second pre-training assessment as the reference.

#### Analysis of MEP amplitudes

MEP amplitude was quantified from suprathreshold responses: trials delivered at 110% and 120% of the preliminary RMT and at maximum stimulator output. Be-cause MEP amplitudes are right-skewed, peak-to-peak amplitudes were log_10_-transformed before averaging, consistent with the log-space input–output curve fitting. For each participant, muscle and assessment, log amplitudes were first averaged within each intensity *×* pulse-width cell, and the resulting twelve cells (three intensities *×* four pulse widths) were then averaged with equal weight, so that differences in the number of retained trials between cells could not bias the composite. The pre-training value was the mean of the two pre-training assessments, computed in log space. Values were back-transformed to mV, yielding a geometric-mean amplitude, and normalised to the group-mean pre-training value for each muscle (see *Normalisation and outlier screening*). This pulse-width-averaged MEP amplitude was the primary measure of corticospinal facilitation, drawing on all four pulse widths as SDTC and rheobase do, and was analysed in the same way as SDTC and rheobase: a 2 *×* 2 (Time *×* Muscle) repeated-measures ANOVA comparing pre-training with the mean of the three post-training assessments, followed by paired *t*-tests in each muscle, and a 2 *×* 4 ANOVA including all four time points. To test whether training effects depended on pulse width, the same aggregation was repeated separately for each pulse width (equal-weight mean of the three intensity cells, normalised per muscle and pulse width) and analysed with a Time *×*

Muscle *×* Pulse-width repeated-measures ANOVA; paired *t*-tests compared each post-training as- sessment against baseline for each muscle and pulse width, with the three within-muscle comparisons Holm–Bonferroni corrected.

#### Resting motor threshold and forward strength–duration modelling

Percent change in RMT_I–O_ from baseline was analysed using a three-way repeated-measures ANOVA with within-subject factors Time (5, 30, 60 min), Muscle (trained, untrained) and Pulse Width (30, 60, 90, 120 *µ*s; Supplementary Table 3). As no effects involving Time were observed, change scores were averaged across the three post-training blocks and analysed using a reduced two-way ANOVA (Muscle *×* Pulse Width), followed by one-sample *t*-tests against zero for each muscle and pulse width. To assess whether the observed SDTC and rheobase changes were of sufficient magnitude to produce detectable threshold shifts, we used a forward strength–duration model in which the strength–duration membrane model was parameterised with each participant’s pre and post-training SDTC and rheobase estimates and used to compute predicted RMT at each of the four pulse widths. Predicted percent change in RMT from baseline was then computed at each pulse width.

#### Brain–behaviour correlations

Associations between training-induced neurophysiological change and behavioural learning were quantified using Spearman rank correlations. Spearman’s *ρ* was used throughout because we did not have strong a priori grounds to expect linear relationships between neurophysiological change scores and behavioural outcomes, and rank-based estimators are more robust to the influence of individual points in modest samples. For each participant and muscle, neurophysiological change scores were computed as the difference between pooled post-training (mean of 5 min, 30 min, 60 min) and pre (mean of the two pre-training blocks) values for SDTC, rheobase, and MEP amplitude. Two primary behavioural outcomes were tested: slope of the training curve, indexing the rate of motor acquisition during training, and 1 h retention, indexing the persistence of training gains beyond the practice context; 24 h retention values (with and without feedback; *N* = 17) were treated as supplementary outcomes given the small sample size. The primary brain–behaviour family, tested per muscle and per behavioural outcome, comprised three correlations: SDTC, rheobase, and MEP amplitude averaged across the four pulse widths. As a sensitivity analysis, MEP amplitude at each individual pulse width (30, 60, 90 and 120 *µ*s) was tested against each outcome in the trained muscle. All correlation families were corrected using the Benjamini–Hochberg FDR procedure.

To test whether intrinsic excitability changes predicted behavioural learning independently of corticospinal facilitation, Spearman partial correlations were computed between each intrinsic excitability measure (SDTC, rheobase) and each behavioural outcome, controlling for the change in pulse-width-averaged MEP amplitude. Reverse partial correlations (MEP amplitude versus each behavioural outcome, controlling for SDTC or rheobase) were also computed. These tests serve to decompose associations identified in the primary analyses; Benjamini–Hochberg correction was applied within each muscle across the four partial correlations for each behavioural outcome. Sensitivity analyses repeated the partial correlations in the trained muscle using MEP amplitude at each individual pulse width (30, 60, 90 or 120 *µ*s) as the covariate. Separately, the direct relationship between SDTC or rheobase change and MEP change within each muscle was also examined using Spearman correlations.

The robustness of all correlations was assessed using leave-one-out (LOO) influence analysis. For each correlation, each participant was removed in turn and the coefficient recomputed and the range of LOO coefficients is reported. Scatter plots display Theil–Sen robust regression lines with bootstrap 95% confidence intervals (10,000 resamples).

*SDTC–rheobase correlations.* Because SDTC and rheobase are jointly derived from the same strength–duration fit, a strong correlation between them is structurally expected. To test whether training altered the slope of this relationship, and whether any such change was specific to the trained muscle, we fitted a linear mixed-effects (LME) model with SDTC as the dependent variable, using one pre-training value (mean of the pre-training assessments) and one post-training value (mean of the post-training assessments) per participant and muscle. Fixed effects were Muscle (categorical: trained, untrained; untrained as reference), Time (categorical: pre, post; pre as reference), grand-mean-centred rheobase (continuous), and all interactions up to and including the three-way Rheobase *×* Time *×* Muscle term. Grand-mean centring of rheobase improves the interpretability of main effects and interaction coefficients while retaining both within- and between-subject variance. Random effects included a by-participant random intercept and a by-participant random slope for rheobase, accounting for individual differences in both pre SDTC and the strength of the SDTC–rheobase relationship. Models were fitted by maximum likelihood.

Muscle specificity of the training-induced slope change was tested with a likelihood-ratio test comparing the full model against a reduced model without the three-way interaction term. Per-muscle pre-to-post slope changes (Δ*β*) were then extracted as follow-up contrasts from the full model: for the untrained muscle this corresponded directly to the Rheobase *×* Time coefficient, while for the trained muscle it was obtained by summing the Rheobase *×* Time and three-way interaction coefficients, with the standard error derived from the full coefficient covariance matrix and evaluated against a *t* distribution with the model’s residual degrees of freedom.

### Computational modelling

#### Model architecture

Because SDTC and rheobase are properties of the axonal membrane, we used a published myelinated axon model whose nodal conductances had been fitted to axonal excitability data: the McIntyre–Richardson–Grill (MRG) double-cable fibre model83, in the small-calibre refit of Peña, Pelot and Grill35. All parameters were used as published. The model is a peripheral fibre model and carries no cortex-specific conductances.

Simulations were run in the NEURON simulation environment (v9.0.1)84 through the PyFibers package (v0.8.5)85. A straight fibre of 3.66 *µ*m diameter (internodal axon diameter 2.0 *µ*m) was built with 25 nodes of Ranvier, giving 265 compartments over a total length of 8.89 mm with a node-to-node distance of 370.6 *µ*m. Nodes were 1.0 *µ*m long with an axon diameter of 1.012 *µ*m; each internode comprised paranodal (MYSA, 3.0 *µ*m; FLUT, 20.5 *µ*m) and internodal (STIN, 53.8 *µ*m) segments arranged symmetrically about its midpoint, with an axon diameter of 2.000 *µ*m beneath the myelin (g-ratio 0.546, 33 lamellae). All geometry followed the model’s diameter-dependent polynomials. Active conductances were present at nodes of Ranvier only: transient sodium 2.333 S/cm2, persistent sodium 0.01 S/cm2, slow potassium 0.1156 S/cm2 and leak 0.007 S/cm2, with reversal potentials +50 mV (Na+) and *−*90 mV (K+ and leak), nodal capacitance 2.0 *µ*F/cm2 and axial resistivity 70 Ω*·*cm. Internodal compartments were passive (reversal *−*80 mV; conductance 2.77 *×* 10*^−^*^4^ S/cm2 at MYSA and 5.46 *×* 10*^−^*^5^ S/cm2 at FLUT and STIN), with the myelin sheath represented as the second cable of the double-cable formulation (1.52 *×* 10*^−^*^5^ S/cm2, 1.52 *×* 10*^−^*^3^ *µ*F/cm2). Kinetics were temperature-corrected to 37*◦*C, and the two terminal nodes were passive (package default), leaving 23 active nodes.

Every simulation was preceded by equilibration from *−*200 ms to *t* = 0. The unmodified fibre settled to *−*80.87 mV at the central node (*−*80.78 mV at the node of action potential initiation) with a residual drift of 6.6 *×* 10*^−^*^5^ mV over a further 5 ms; leak and slow potassium accounted for 60% and 36% of the nodal resting conductance, the transient sodium window current 4.0% and persistent sodium 0.6%. Because a change in any nodal conductance can itself shift the resting potential, the settled potential and its stability were measured for every condition. Conditions were accepted only if the membrane potential drifted by less than 0.5 mV over 5 ms with no spontaneous action potentials, which distinguishes a stable but tonically depolarised resting state from a loss of resting stability.

#### Stimulation and threshold determination

The model was stimulated with the measured cTMS electric-field waveforms of the pulses used in the experimental protocol (30, 60, 90 and 120 *µ*s), recorded at 2 MHz and applied as a spatially uniform extracellular field oriented along the axon, with the stimulus amplitude expressed as the peak field in V/m. Because the field is uniform and the axon straight, the activating function is zero everywhere except at the sealed ends; action potentials therefore initiated at the last active node at the depolarised end (node 24 of 25) and propagated along the fibre. This follows from the uniform-field idealisation rather than from the fibre model.

Simulations used a fixed 0.5 *µ*s time step — equal to the sampling interval of the recorded waveform, so that the model was driven by the measured record without decimation or interpolation — and ran for 2 ms with pulse onset at 0.5 ms. The activation threshold at each pulse width was determined by bisection on the field amplitude, terminated at a relative resolution of 10*^−^*^5^, with an action potential defined as a crossing of *−*30 mV at the midpoint of the central node. SDTC and rheobase were then estimated from the four simulated thresholds using the same leaky-integrator membrane model and fitting procedure applied to the experimental data (see *SDTC and rheobase estimation*).

Because model thresholds are electric-field amplitudes, the fitted rheobase is expressed in V/m and is not directly comparable to the experimental rheobase in % MSO; all model results are therefore reported as percentage changes from the model’s own baseline.

#### Parameter sensitivity analyses

Five parameters were varied independently, each across 11 values with the baseline value at the centre of the grid. Ranges differ between parameters, each chosen so that its sweep spans changes in SDTC and rheobase of the order observed experimentally. Transient sodium (*g*NaT) was scaled over 0.85–1.15*×* baseline, persistent sodium (*g*NaP) was scaled over 0.25–4*×* and slow potassium (*g*K) over 0.5–2*×*. Resting membrane potential (*V*_m_) was varied over *±*1 mV in 0.2 mV steps by shifting the nodal and internodal leak reversal potentials together, with the required shift solved numerically for each target and the sweep defined by the achieved change in settled potential at the node of action potential initiation (converged to within 10*^−^*^4^ mV). Axon diameter, defined as the internodal axon diameter beneath the myelin, was varied linearly over 1–3 *µ*m (0.5–1.5*×*; fibre diameters 1.85–5.47 *µ*m), with every geometric parameter rescaled by the model’s diameter polynomials, so that total fibre length varied between 3.2 and 14.0 mm; this range spans the reported calibre of major intracortical axon collaterals in motor cortex38 and lies within the model’s validity range. At every value the full strength–duration protocol was repeated and SDTC and rheobase were re-estimated; every condition met the resting-stability criterion above. For each parameter we then identified the value that reproduced the median within-subject rheobase change in the trained muscle (*−*4.9%; pre-training versus the mean of the three post-training assessments) and recorded the accompanying change in SDTC, bracketing it with the adjacent pair of simulated points from that parameter’s sweep and refining by safeguarded false position (Illinois) until the simulated rheobase change was within 0.003 percentage points of *−*4.9%. The median was used as the matching target because it is less sensitive than the mean to the skewed distribution of individual change scores.

## Funding

B.C. and R.H.’s contribution was supported by the BBSRC (UKRI5216) and the Academy of Medical Sciences Springboard scheme (SBF009\1073), which is funded by British Heart Foundation, Diabetes UK, the Government Department for Science, Innovation and Technology (DSIT) and Wellcome. A.V.P.’s contribution to the reported research was supported by the National Institutes of Health under Award Numbers RF1MH124943, R01NS117405, and R01MH128422. The content is solely the responsibility of the authors and does not necessarily represent the official views of the funding agencies.

## Acknowledgements

We would like to thank Mathew Yarossi and Gene Tunik for supplying the Synergy software for cTMS control and EMG signal acquisition, Andy Palmer for supplying the custom-built accelerometer, and Brainbox Ltd. for the use of the TMS device.

## Competing interests

A.V.P. is an inventor on patents on TMS technology and has received: consulting fees and patent royalties for a license on the cTMS (Elevate TMS) technology used in this study from Rogue Research; equity options, scientific advisory board membership, research funding and consulting fees from Ampa Health; equity options, consulting fees and travel support from Magnetic Tides; equipment loan from MagVenture; hardware donation from Magstim; and research funding from Motif. The other authors have no relevant disclosures.

## Author contributions

Benjamin Clennell: Conception or design of the work; Acquisition, analysis, or interpretation of data for the work; drafting the work or revising it critically for important intellectual content. Binyu Luo: Acquisition, analysis or interpretation of data for the work. Kei Tung Lam: Acquisition, analysis or interpretation of data for the work. Yu Hu: Acquisition, analysis or interpretation of data for the work. Angel V. Peterchev: Acquisition, analysis, or interpretation of data for the work; drafting the work or revising it critically for important intellectual content. Ricci Hannah: Conception or design of the work; acquisition, analysis, or interpretation of data for the work; drafting the work or revising it critically for important intellectual content. All authors approved the final version of the manuscript, and all agree to be accountable for all aspects of the work in ensuring that questions related to the accuracy or integrity of any part of the work are appropriately investigated and resolved. All persons designated as authors qualify for authorship, and all those who qualify for authorship are listed.

## Data availability

All raw and processed data supporting the findings of this study, including source data for all figures and Supplementary Tables, are available in the King’s College London Research Data Repository at https://doi.org/10.18742/34807857 (CC BY 4.0).

## Code availability

All MATLAB analysis code and the Python axon model code used to generate the results, figures and Supplementary Tables are available in the King’s College London Research Data Repository at https://doi.org/10.18742/34922793 (MIT licence).

## Extended Data Figures

**Extended Data Fig. 1.**
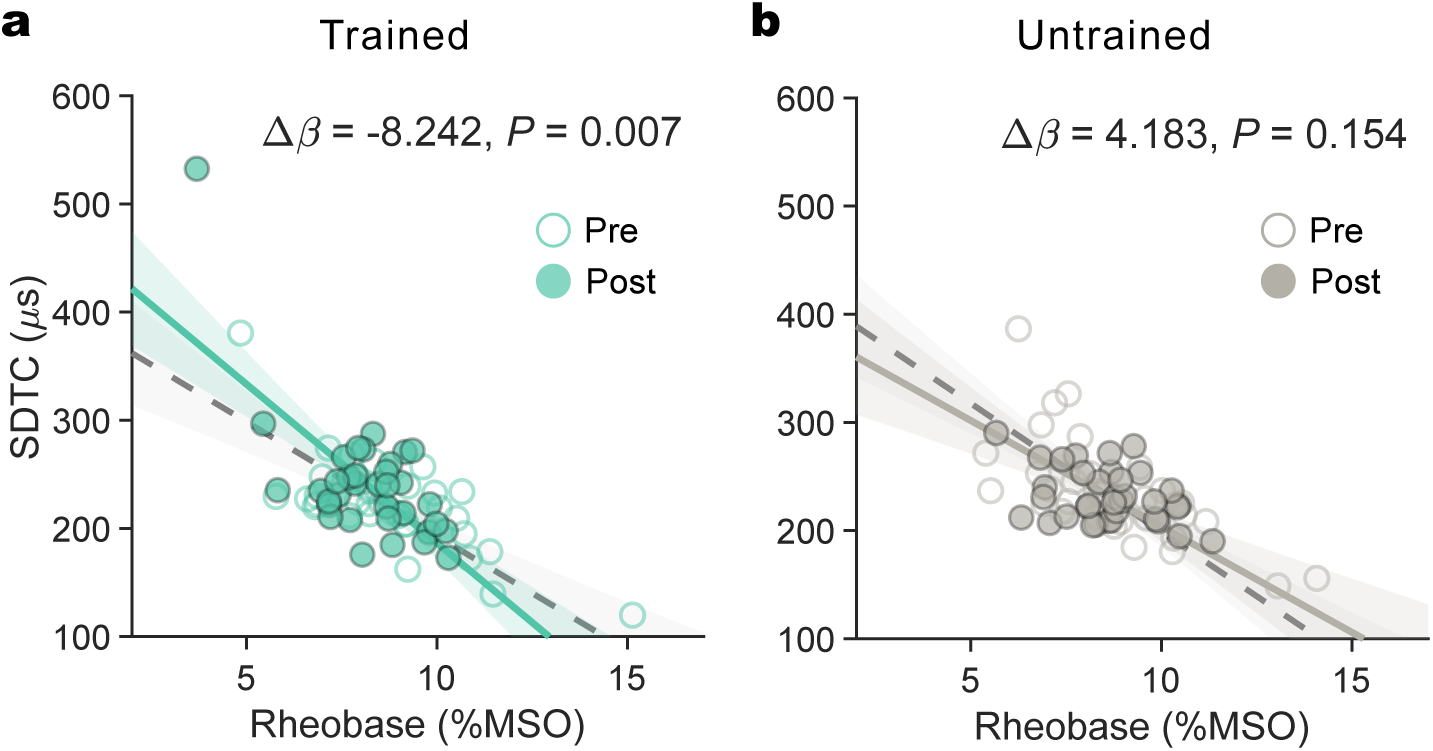
Motor training selectively reshapes the relationship between SDTC and rheobase in the trained muscle. SDTC plotted against rheobase before (Pre, open circles) and after training (Post, filled circles) in the (a) trained and (b) untrained muscles. Pre-training values are the mean of the two pre-training assessments; post-training values are the mean of the 5, 30 and 60 min assessments. Lines denote the fixed-effect SDTC–rheobase slopes from the linear mixed-effects model at each time point. Training altered the SDTC–rheobase slope differently in the two muscles (Rheobase *×* Time *×* Muscle, likelihood-ratio test: *χ*^2^(1) = 9.17, *P* = 0.002). The negative slope steepened after training in the trained muscle (Δ*β* = *−*8.24, *P* = 0.007) but not in the untrained muscle (Δ*β* = 4.18, *P* = 0.15). Δ*β* denotes the pre-to-post change in slope (*µ*s per %MSO), estimated within each muscle as a follow-up contrast from the full model. *N* = 35.

**Extended Data Fig. 2.**
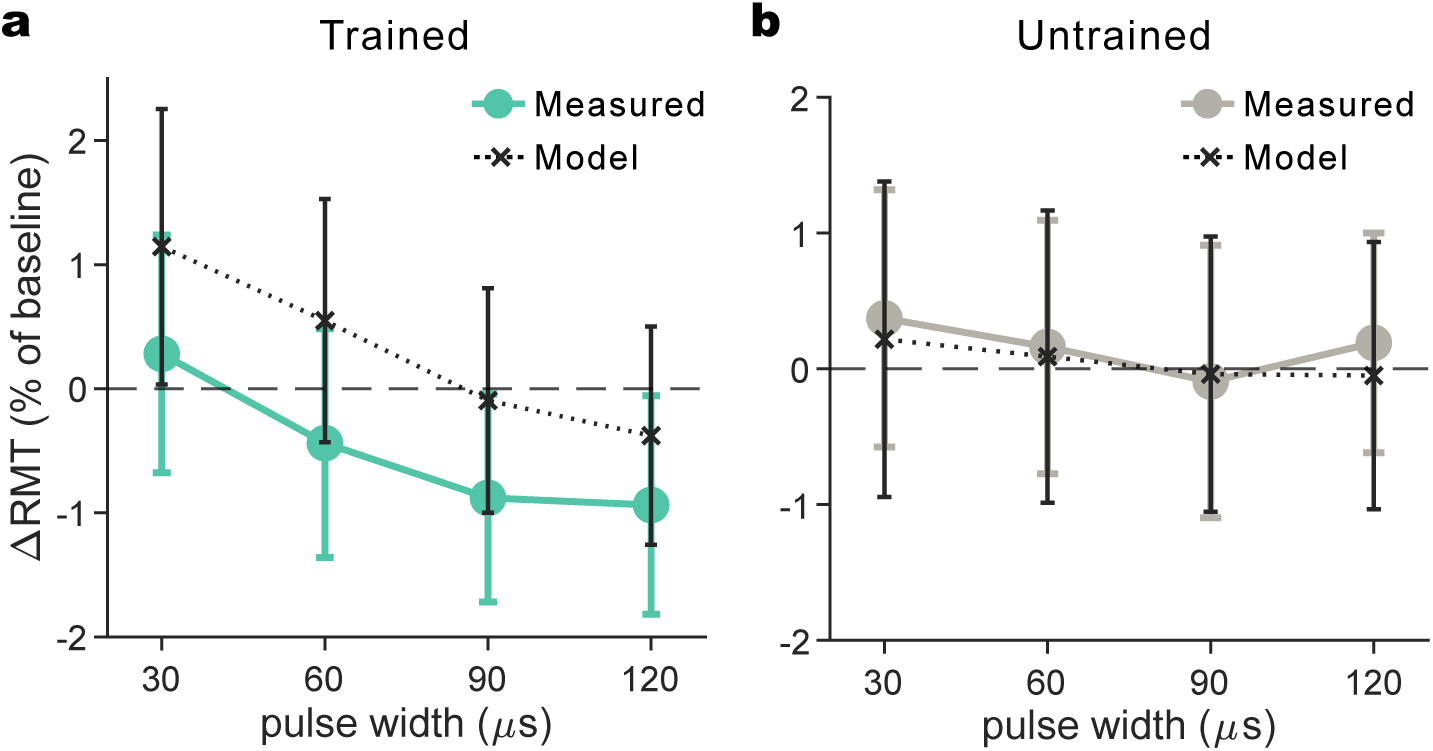
Resting motor threshold was unchanged after training, consistent with forward model predictions. Measured percent change in resting motor threshold (ΔRMT) from baseline across the four pulse widths in the (a) trained and (b) untrained muscles, averaged across the three post-training timepoints, shown alongside predictions of the forward strength–duration model. A two-way repeated-measures ANOVA on the post-averaged data (Muscle *×* Pulse Width) revealed no main effect of Muscle (*F* (1, 34) = 1.93, *P* = .174, *η*^2^_*p*_ = .05), no main effect of Pulse Width (*F* (3, 102) = 1.40, *P*_GG_ = .252, *η*^2^_*p*_ = .04), and no Muscle *×* Pulse Width interaction (*F* (3, 102) = 1.38, *P*_GG_ = .255, *η*^2^_*p*_ = .04). One-sample tests against zero confirmed that RMT did not change in either muscle at any pulse width (all *|d_z_| ≤* 0.18, all *P ≥ .*29; uncorrected). The full Time *×* Muscle *×* Pulse Width analysis, which revealed no effects involving Time, is reported in Supplementary Table 3. For the model, the leaky-integrator membrane model was parameterised with each participant’s empirical pre- and post-training SDTC and rheobase estimates to compute a predicted RMT at each pulse width. The model predicts sub-percent changes in RMT, producing only a slight negative gradient at longer pulse widths in the trained muscle. The observed RMT data closely track these predictions. Data are mean *±* SEM.

**Extended Data Fig. 3.**
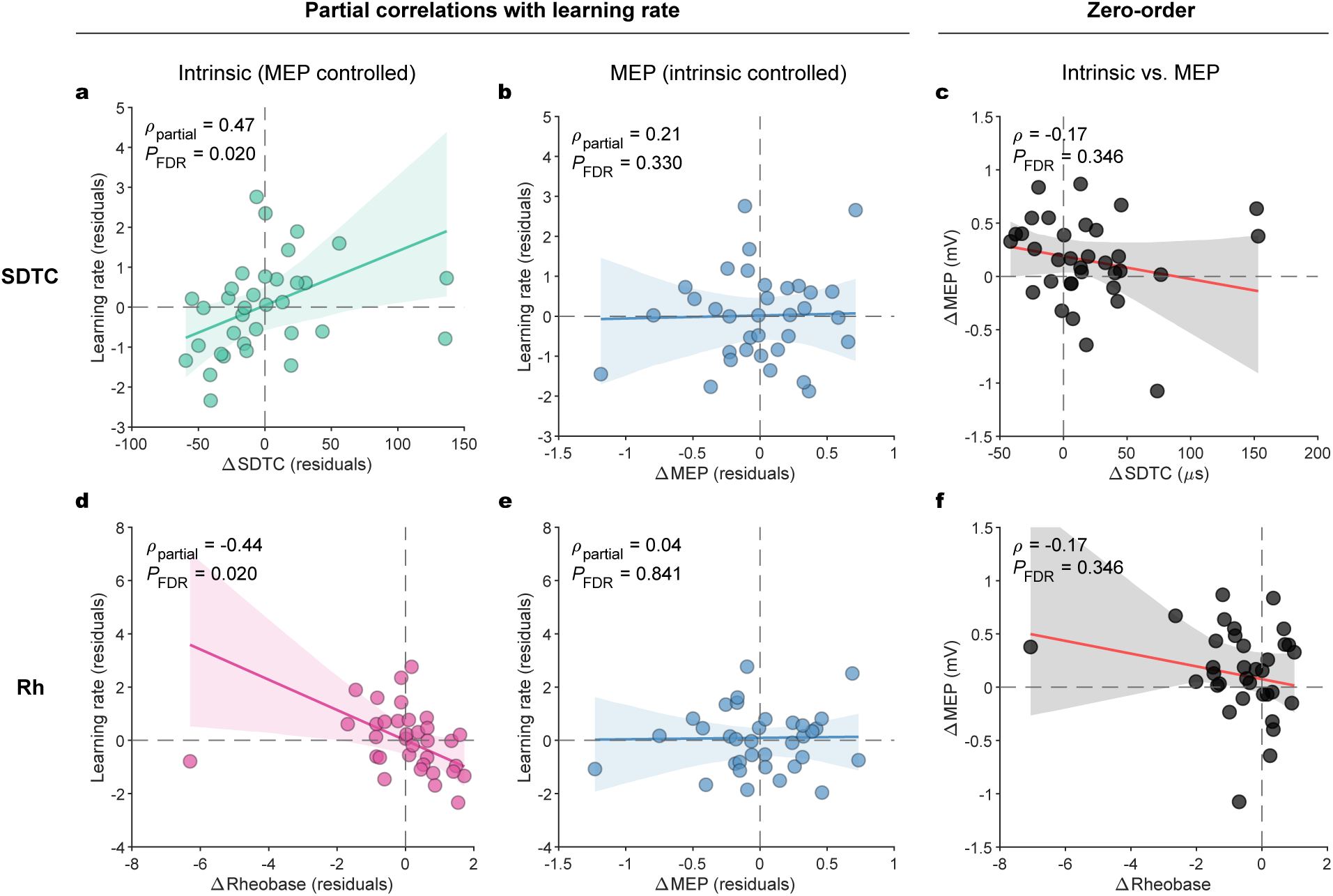
Intrinsic excitability changes predict learning independently of corticospinal facilitation. Partial and zero-order Spearman correlations in the trained muscle, showing the dissociation between intrinsic excitability measures (SDTC, top row; rheobase, bottom row) and MEP amplitude changes. (a) SDTC change predicts learning rate after controlling for concurrent MEP change. (b) Relationship between MEP change and learning rate remains non-significant when controlling for SDTC change. (c) SDTC change and MEP change are not significantly correlated. (d) rheobase change predicts learning rate after controlling for concurrent MEP change. (e) Relationship between MEP change and learning rate remains non-significant when controlling for rheobase change. (f) Rheobase change and MEP change are not significantly correlated. (a,b,d,e) display residuals from the partial correlation procedure; (c,f) display raw change scores with original units. Each dot represents one participant; solid lines denote Theil–Sen robust regression fits and shaded bands the 95% bootstrap confidence interval. *P*_FDR_, Benjamini–Hochberg-corrected within each family: the four partial correlations (a,b,d,e) and the two intrinsic–MEP correlations (c,f); *N* = 34. Results were equivalent regardless of the pulse width used for the covariate MEP change (Supplementary Table 8).

**Extended Data Fig. 4.**
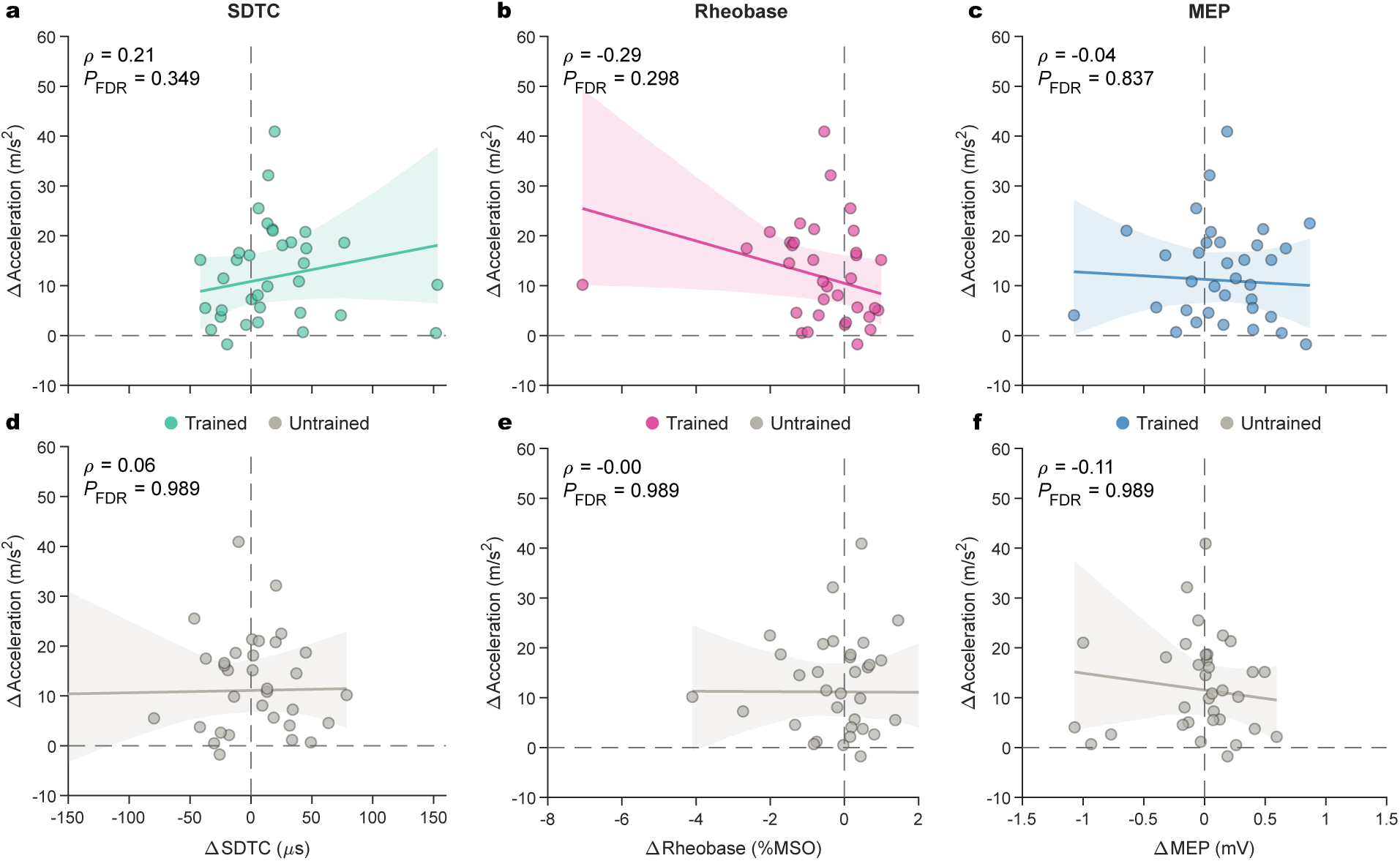
Associations between training-induced neurophysiological changes and short-term retention. Spearman correlations between pre-to-post change in each neurophysiological measure (collapsed across the three post-training time points) and 1-hour retention (change in peak acceleration from baseline), shown separately for the trained muscle (a–c) and untrained control muscle (d–f). No neurophysiological measure in the trained or untrained muscle was significantly associated with retention. Each dot represents one participant (*N* = 34); solid lines denote the Theil–Sen robust regression fit and shaded bands the 95% bootstrap confidence interval.

## Notes

https://doi.org/10.18742/34807857

https://doi.org/10.18742/34922793

