## Supplementary Information for "Motor learning engages intrinsic plasticity in the human cortex"

#### Intrinsic excitability (SDTC and rheobase)

##### Full ANOVA results and sensitivity analyses (Tables S1–S2)

The main text reports the pre-versus-post ($2 \times 2$) and time-course ($2 \times 4$) Muscle × Time ANOVAs for SDTC and rheobase. Tables S1 and S2 give the full results, alongside two sensitivity analyses. First, baseline SDTC and rheobase were stable across the two pre-training assessments in the trained muscle but differed in the untrained muscle. This raised the possibility that the muscle-specific effects reflected drift between baseline assessments rather than training, so we re-referenced change scores to the immediate pre-training assessment alone (Panel B). Second, to test whether the effects depended on extreme values, we excluded four participants whose SDTC or rheobase change exceeded 3.5× the scaled median absolute deviation in either muscle (Panel C; $N = 31$). Under both sensitivity specifications the results were largely unchanged, except for a slight attenuation of the Muscle main effect in the SDTC $2 \times 4$ ANOVA after outlier exclusion ($P = .055$).

**Table S1: ANOVA results for strength–duration time constant (SDTC).**

| Analysis | Effect | $F$ | df | $P$ | $P_{GG}$ | $\eta_p^2$ |
|---|---|---|---|---|---|---|
| *Panel A. Primary analysis ($N = 35$)* | | | | | | |
| Pre vs post ($2 \times 2$) | Time | 1.69 | 1, 34 | .202 | – | .047 |
| | Muscle | 2.40 | 1, 34 | .130 | – | .066 |
| | Time × Muscle | **7.06** | 1, 34 | **.012** | – | .172 |
| Time course ($2 \times 4$) | Time | 0.63 | 3, 102 | .595 | .557 | .018 |
| | Muscle | **4.37** | 1, 34 | **.044** | – | .114 |
| | Time × Muscle | 1.38 | 3, 102 | .254 | .259 | .039 |
| *Panel B. Sensitivity: single pre-training baseline ($N = 35$)* | | | | | | |
| Pre vs post ($2 \times 2$) | Time | 0.04 | 1, 34 | .836 | – | .001 |
| | Muscle | 3.51 | 1, 34 | .070 | – | .093 |
| | Time × Muscle | **9.15** | 1, 34 | **.005** | – | .212 |
| Time course ($2 \times 4$) | Time | 0.21 | 3, 102 | .891 | .854 | .006 |
| | Muscle | **8.29** | 1, 34 | **.007** | – | .196 |
| | Time × Muscle | 2.51 | 3, 102 | .063 | .086 | .069 |
| *Panel C. Sensitivity: MAD outlier exclusion ($N = 31$)* | | | | | | |
| Pre vs post ($2 \times 2$) | Time | 1.63 | 1, 30 | .211 | – | .052 |
| | Muscle | 1.72 | 1, 30 | .199 | – | .054 |
| | Time × Muscle | **4.88** | 1, 30 | **.035** | – | .140 |
| Time course ($2 \times 4$) | Time | 0.40 | 3, 90 | .753 | .706 | .013 |
| | Muscle | 3.98 | 1, 30 | .055 | – | .117 |
| | Time × Muscle | 1.20 | 3, 90 | .313 | .309 | .039 |

*Note:* Analysis: pre vs post, Time (pre, post; post is the mean of the 5, 30 and 60 min assessments) × Muscle; time course, Time (pre, 5, 30, 60 min) × Muscle. Panel A, pre-training reference is the mean of the two pre-training assessments; Panel B, immediate pre-training assessment only; Panel C, exclusion of four participants exceeding 3.5× the scaled median absolute deviation on SDTC or rheobase change in either muscle, with data re-normalised to the reduced sample's grand-mean baseline. $P_{GG}$, Greenhouse–Geisser-corrected $P$ (not applicable to single-df effects); $\eta_p^2$, partial eta-squared. Bold, $P < .05$.

#### Table S2: ANOVA results for rheobase.

| Analysis | Effect | $F$ | df | $P$ | $P_{GG}$ | $\eta_p^2$ |
|---|---|---|---|---|---|---|
| *Panel A. Primary analysis ($N = 35$)* | | | | | | |
| Pre vs post ($2 \times 2$) | Time | 2.80 | 1, 34 | .103 | – | .076 |
| | Muscle | 2.91 | 1, 34 | .097 | – | .079 |
| | Time $\times$ Muscle | **6.51** | 1, 34 | **.015** | – | .161 |
| Time course ($2 \times 4$) | Time | 1.26 | 3, 102 | .291 | .291 | .036 |
| | Muscle | **6.95** | 1, 34 | **.013** | – | .170 |
| | Time $\times$ Muscle | 1.39 | 3, 102 | .251 | .254 | .039 |
| *Panel B. Sensitivity: single pre-training baseline ($N = 35$)* | | | | | | |
| Pre vs post ($2 \times 2$) | Time | 0.02 | 1, 34 | .876 | – | .001 |
| | Muscle | **6.39** | 1, 34 | **.016** | – | .158 |
| | Time $\times$ Muscle | **13.35** | 1, 34 | **<.001** | – | .282 |
| Time course ($2 \times 4$) | Time | 0.16 | 3, 102 | .921 | .892 | .005 |
| | Muscle | **18.85** | 1, 34 | **<.001** | – | .357 |
| | Time $\times$ Muscle | **3.75** | 3, 102 | **.013** | **.016** | .099 |
| *Panel C. Sensitivity: MAD outlier exclusion ($N = 31$)* | | | | | | |
| Pre vs post ($2 \times 2$) | Time | 2.01 | 1, 30 | .167 | – | .063 |
| | Muscle | 4.00 | 1, 30 | .054 | – | .118 |
| | Time $\times$ Muscle | **6.44** | 1, 30 | **.017** | – | .177 |
| Time course ($2 \times 4$) | Time | 0.59 | 3, 90 | .624 | .594 | .019 |
| | Muscle | **8.01** | 1, 30 | **.008** | – | .211 |
| | Time $\times$ Muscle | 1.42 | 3, 90 | .243 | .247 | .045 |

*Note:* Repeated-measures ANOVAs on normalised rheobase; analyses, panels and abbreviations as in Table S1. Bold, $P < .05$.

#### Sensitivity of intrinsic excitability changes to MEP criterion (Fig. S1)

Our primary analysis estimated motor thresholds using a 200 $\mu$V MEP criterion. To test whether the training-induced changes in SDTC and rheobase depended on this choice, we repeated the analysis using thresholds derived at 50, 100 and 300 $\mu$V, alongside the original 200 $\mu$V criterion. Lower criteria lie closer to the point of activation but fall at the lower edge of the sampled intensity range, whereas higher criteria are constrained by the pulse-width-specific maximum stimulator output. At each criterion, thresholds were taken from the same fitted Boltzmann functions and SDTC and rheobase were re-estimated. Their pre-to-post change was entered into a Criterion × Muscle repeated-measures ANOVA ($N = 35$). Neither term approached significance for SDTC (Criterion: $F(3, 102) = 0.05$, $P_{GG} = 0.86$, $\eta_p^2 = 0.002$; Criterion × Muscle: $F(3, 102) = 0.15$, $P_{GG} = 0.75$, $\eta_p^2 = 0.004$) or rheobase (Criterion: $F(3, 102) = 0.32$, $P_{GG} = 0.57$, $\eta_p^2 = 0.009$; Criterion × Muscle: $F(3, 102) = 0.96$, $P_{GG} = 0.34$, $\eta_p^2 = 0.028$). The training-related change in intrinsic excitability therefore did not depend on the criterion used to define threshold (Supplementary Fig. S1).

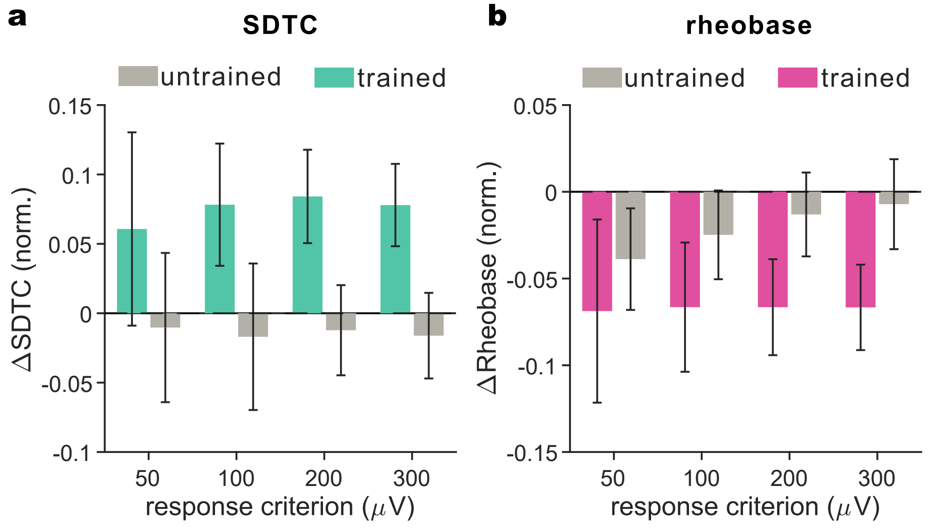

**Figure S1: Training-related changes in SDTC and rheobase are independent of the response criterion used to derive motor threshold.** Pre-to-post change in (a) SDTC and (b) rheobase in the trained (coloured) and untrained control (grey) muscles, with resting motor threshold derived from the same fitted input–output curves at four response criteria (50, 100, 200 and 300 $\mu$V). Change scores are the collapsed post-training value minus baseline, with each criterion normalised to its own group grand-mean baseline (see Methods), so that values are expressed as a proportion of baseline. Changes were in the same direction and of similar magnitude across all four criteria in both muscles, with no main effect of criterion and no Criterion × Muscle interaction for either measure. Data are mean ± SEM; $N = 35$.

#### Resting motor threshold (Table S3)

Extended Data Figure 2 reports the change in resting motor threshold after training. To test whether threshold changed at any post-training assessment, in either muscle or at any pulse width, we analysed percent change in RMT from baseline with a full Time × Muscle × Pulse Width repeated-measures ANOVA (Table S3). No effect reached significance, including all terms involving Time. RMT change was therefore averaged across the three post-training assessments in Extended Data Figure 2.

**Table S3: Full three-way analysis of resting motor threshold change.**

| Effect | $F$ | df | $P$ | $P_{\mathrm{GG}}$ | $\eta_p^2$ |
|---|---|---|---|---|---|
| Time | 0.19 | 2, 68 | .824 | .761 | .006 |
| Muscle | 1.93 | 1, 34 | .174 | – | .054 |
| PulseWidth | 1.40 | 3, 102 | .247 | .252 | .040 |
| Time × Muscle | 0.81 | 2, 68 | .448 | .445 | .023 |
| Time × PulseWidth | 1.09 | 6, 204 | .367 | .364 | .031 |
| Muscle × PulseWidth | 1.38 | 3, 102 | .252 | .255 | .039 |
| Time × Muscle × PulseWidth | 0.13 | 6, 204 | .992 | .968 | .004 |

*Note:* $P_{\mathrm{GG}}$, Greenhouse–Geisser-corrected $P$; $\eta_p^2$, partial eta-squared.

### MEP amplitude

#### Full ANOVA results (Table S4)

The main text reports MEP amplitude averaged across the four pulse widths. Table S4 gives the full ANOVA results and additionally tests whether facilitation differed between pulse widths. Training altered MEP amplitude differently in the trained and untrained muscles, with a significant Time × Muscle interaction for pulse-width-averaged MEP amplitude in both the pre-versus-post and time-course analyses (Table S4, Panels A and B). In the analysis by pulse width, the Time × Muscle interaction remained significant and did not interact with pulse width, providing no evidence that facilitation differed across pulse widths (Table S4, Panel C).

**Table S4: ANOVA results for MEP amplitude.**

| Effect | $F$ | df | $P$ | $P_{\mathrm{GG}}$ | $\eta_p^2$ |
|---|---|---|---|---|---|
| *Panel A. Pulse-width-averaged MEP, pre vs post (2 × 2)* | | | | | |
| Time | 0.65 | 1, 34 | .427 | – | .019 |
| Muscle | 0.56 | 1, 34 | .458 | – | .016 |
| Time × Muscle | **11.40** | 1, 34 | **.002** | – | .251 |
| *Panel B. Pulse-width-averaged MEP, time course (2 × 4)* | | | | | |
| Time | 0.94 | 3, 102 | .425 | .408 | .027 |
| Muscle | 1.17 | 1, 34 | .287 | – | .033 |
| Time × Muscle | **5.98** | 3, 102 | **<.001** | **.001** | .150 |
| *Panel C. MEP by pulse width (2 × 4 × 4)* | | | | | |
| Time | 1.08 | 3, 102 | .360 | .352 | .031 |
| Muscle | 1.15 | 1, 34 | .291 | – | .033 |
| PulseWidth | 0.09 | 3, 102 | .964 | .947 | .003 |
| Time × Muscle | **6.00** | 3, 102 | **<.001** | **.001** | .150 |
| Time × PulseWidth | 0.81 | 9, 306 | .608 | .550 | .023 |
| Muscle × PulseWidth | 0.30 | 3, 102 | .829 | .805 | .009 |
| Time × Muscle × PulseWidth | 0.50 | 9, 306 | .871 | .807 | .015 |

*Note:* Panel A, Time (pre, post; post is the mean of the 5, 30 and 60 min assessments) × Muscle, on MEP amplitude averaged across pulse widths. Panel B, Time (pre, 5, 30, 60 min) × Muscle on the same measure. Panel C, Time × Muscle × Pulse width (30, 60, 90, 120 $\mu$s). $P_{\mathrm{GG}}$, Greenhouse–Geisser-corrected $P$ (not applicable to single-df effects); $\eta_p^2$, partial eta-squared. Bold, $P < .05$.

#### Sensitivity to outlier exclusion (Table S5)

As for SDTC and rheobase, we tested whether the muscle-specific MEP facilitation depended on extreme responders: we repeated the pulse-width-averaged analyses after excluding four participants whose MEP change exceeded 3.5× the scaled median absolute deviation in either muscle ($N = 31$). All four showed large decreases, predominantly in the untrained muscle. The Muscle × Time interaction was preserved in both the pre-versus-post and time-course analyses (Table S5).

**Table S5: Sensitivity analysis of MEP amplitude effects.**

| Effect | $F$ | df | $P$ | $P_{\mathrm{GG}}$ | $\eta_p^2$ |
|---|---|---|---|---|---|
| *Panel A. Pulse-width-averaged MEP, pre vs post (2 × 2)* | | | | | |
| Time | **14.53** | 1, 30 | **<.001** | – | .326 |
| Muscle | 0.22 | 1, 30 | .643 | – | .007 |
| Time × Muscle | **5.59** | 1, 30 | **.025** | – | .157 |
| *Panel B. Pulse-width-averaged MEP, time course (2 × 4)* | | | | | |
| Time | **5.54** | 3, 90 | **.002** | **.003** | .156 |
| Muscle | 0.45 | 1, 30 | .508 | – | .015 |
| Time × Muscle | **3.86** | 3, 90 | **.012** | **.016** | .114 |

*Note:* Repeated-measures ANOVAs on normalised, pulse-width-averaged MEP amplitude after excluding four participants exceeding 3.5× the scaled median absolute deviation on MEP change in either muscle ($N = 31$). Data were re-normalised to the reduced sample's grand-mean baseline. Panel A, Time (pre, post; post is the mean of the 5, 30 and 60 min assessments) × Muscle. Panel B, Time (pre, 5, 30, 60 min) × Muscle. $P_{\mathrm{GG}}$, Greenhouse–Geisser-corrected $P$ (not applicable to single-df effects); $\eta_p^2$, partial eta-squared. Bold, $P < .05$.

### Brain–behaviour associations

#### Primary correlations (Table S6)

Figure 4 and Extended Data Figs 3 and 4 report the primary brain–behaviour correlations. Table S6 gives the full results for both muscles, with bootstrap confidence intervals and leave-one-out ranges. Partial correlations isolate each intrinsic measure from concurrent corticospinal facilitation, and vice versa, and are reported only for the learning rate where significant zero-order correlations were found.

**Table S6: Primary brain–behaviour correlations.**

| Analysis Type | Variable | $\rho$ | 95% CI | $P$ | $P_{\text{FDR}}$ | LOO Range |
|---|---|---|---|---|---|---|
| **Panel A. Learning Rate** | | | | | | |
| *Trained Muscle (APB)* | | | | | | |
| Zero-order | ΔSDTC | **.440** | [0.102, 0.705] | **.010** | **.015** | [0.40, 0.51] |
| | ΔRheobase | **−.452** | [−0.672, −0.154] | **.008** | **.015** | [−0.51, −0.42] |
| | ΔMEP | .110 | [−0.249, 0.454] | .534 | .534 | [0.03, 0.18] |
| Partial | ΔSDTC \| ΔMEP | **.468** | [0.152, 0.713] | **.006** | **.020** | [0.430, 0.541] |
| | ΔRheobase \| ΔMEP | **−.443** | [−0.676, −0.108] | **.010** | **.020** | [−0.502, −0.393] |
| | ΔMEP \| ΔSDTC | .207 | [−0.164, 0.525] | .248 | .330 | [0.130, 0.268] |
| | ΔMEP \| ΔRheobase | .036 | [−0.324, 0.399] | .841 | .841 | [−0.042, 0.093] |
| *Untrained Muscle (ADM)* | | | | | | |
| Zero-order | ΔSDTC | .034 | [−0.295, 0.364] | .849 | .963 | [−0.04, 0.11] |
| | ΔRheobase | −.105 | [−0.406, 0.203] | .552 | .963 | [−0.16, −0.04] |
| | ΔMEP | .008 | [−0.362, 0.391] | .963 | .963 | [−0.06, 0.10] |
| Partial | ΔSDTC \| ΔMEP | .036 | [−0.277, 0.351] | .840 | .979 | [−0.024, 0.103] |
| | ΔRheobase \| ΔMEP | −.105 | [−0.423, 0.221] | .561 | .979 | [−0.179, −0.040] |
| | ΔMEP \| ΔSDTC | .016 | [−0.345, 0.378] | .929 | .979 | [−0.048, 0.092] |
| | ΔMEP \| ΔRheobase | −.005 | [−0.385, 0.384] | .979 | .979 | [−0.092, 0.089] |
| **Panel B. Short-Term Retention (1 h Post-Training)** | | | | | | |
| *Trained Muscle (APB)* | | | | | | |
| Zero-order | ΔSDTC | .210 | [−0.154, 0.530] | .233 | .349 | [0.16, 0.31] |
| | ΔRheobase | −.287 | [−0.558, 0.026] | .099 | .298 | [−0.34, −0.24] |
| | ΔMEP | −.037 | [−0.412, 0.341] | .837 | .837 | [−0.11, 0.04] |
| *Untrained Muscle (ADM)* | | | | | | |
| Zero-order | ΔSDTC | .061 | [−0.314, 0.414] | .731 | .989 | [−0.00, 0.14] |
| | ΔRheobase | −.003 | [−0.342, 0.328] | .989 | .989 | [−0.08, 0.06] |
| | ΔMEP | −.113 | [−0.463, 0.279] | .522 | .989 | [−0.19, −0.05] |

*Note:* Spearman correlations ($\rho$) and partial correlations ($\rho_{\text{partial}}$) with the primary learning rate (Panel A; mean within-block OLS slope of peak acceleration), and zero-order correlations with 1 h retention (Panel B; change from baseline). ΔMEP, change in MEP amplitude averaged across pulse widths (30, 60, 90, 120 $\mu$s). Partial correlations control for ΔMEP when evaluating intrinsic measures, or ΔSDTC/ΔRheobase when evaluating corticospinal measures. $P_{\text{FDR}}$, Benjamini–Hochberg-corrected per muscle (zero-order $k = 3$; partials $k = 4$). 95% CIs from 10,000 bootstrap resamples; LOO range, leave-one-out coefficient range. Bold, $P < .05$.

#### Sensitivity to MEP pulse width (Tables S7–S8)

The brain–behaviour analyses used MEP amplitude averaged across the four pulse widths. To test whether the conclusions depended on this choice, we repeated the analyses in the trained muscle with MEP amplitude at each pulse width (30, 60, 90 and 120 $\mu$s) separately. Each pulse width was tested as a predictor of learning rate and 1 h retention (Table S7), and as the covariate in the partial correlations of SDTC and rheobase with learning rate (Table S8).

MEP change at each pulse width was unrelated to both learning rate and 1 h retention, mirroring the pulse-width-averaged result (Table S7).

**Table S7: Sensitivity of zero-order correlations to MEP pulse width.**

| MEP Variable | $\rho$ | 95% CI | $P$ | LOO Range |
|---|---|---|---|---|
| **Panel A. Rate of Acquisition (Learning Slope)** | | | | |
| $\Delta MEP_{30}$ | .008 | [−0.327, 0.355] | .966 | [−0.054, 0.100] |
| $\Delta MEP_{60}$ | .154 | [−0.192, 0.454] | .384 | [0.102, 0.212] |
| $\Delta MEP_{90}$ | .006 | [−0.359, 0.365] | .974 | [−0.081, 0.064] |
| $\Delta MEP_{120}$ | .058 | [−0.301, 0.411] | .746 | [−0.023, 0.122] |
| **Panel B. Short-Term Retention (1 h)** | | | | |
| $\Delta MEP_{30}$ | −.042 | [−0.401, 0.329] | .813 | [−0.107, 0.048] |
| $\Delta MEP_{60}$ | −.020 | [−0.399, 0.359] | .911 | [−0.084, 0.071] |
| $\Delta MEP_{90}$ | −.225 | [−0.588, 0.164] | .199 | [−0.320, −0.164] |
| $\Delta MEP_{120}$ | −.054 | [−0.408, 0.301] | .760 | [−0.131, 0.024] |

*Note:* Zero-order Spearman correlations between ΔMEP at 30, 60, 90 and 120 $\mu$s and the learning rate (Panel A) and 1 h retention (Panel B); trained APB muscle, $N = 34$. 95% CIs from 10,000 bootstrap resamples; LOO range, leave-one-out coefficient range.

The associations of SDTC and rheobase change with learning rate remained significant whichever pulse width was used as the covariate (Table S8).

**Table S8: Sensitivity of partial correlations to MEP covariate pulse width.**

| Covariate Used | Intrinsic Variable | $\rho_{partial}$ | 95% CI | $P$ | LOO Range |
|---|---|---|---|---|---|
| $\Delta MEP_{30}$ | ΔSDTC | **.451** | [0.137, 0.704] | **.008** | [0.415, 0.536] |
| | ΔRheobase | **−.455** | [−0.682, −0.136] | **.008** | [−0.513, −0.407] |
| $\Delta MEP_{60}$ | ΔSDTC | **.462** | [0.135, 0.715] | **.007** | [0.421, 0.528] |
| | ΔRheobase | **−.436** | [−0.669, −0.122] | **.011** | [−0.495, −0.393] |
| $\Delta MEP_{90}$ | ΔSDTC | **.454** | [0.136, 0.703] | **.008** | [0.418, 0.528] |
| | ΔRheobase | **−.453** | [−0.677, −0.137] | **.008** | [−0.511, −0.414] |
| $\Delta MEP_{120}$ | ΔSDTC | **.456** | [0.132, 0.713] | **.008** | [0.419, 0.532] |
| | ΔRheobase | **−.451** | [−0.681, −0.124] | **.008** | [−0.507, −0.402] |

*Note:* Partial Spearman correlations ($\rho_{partial}$) between intrinsic change (ΔSDTC, ΔRheobase) and learning rate, controlling for ΔMEP at 30, 60, 90 or 120 $\mu$s; trained muscle only. $N = 34$. 95% CIs from 10,000 bootstrap resamples; LOO range, leave-one-out coefficient range. Bold, $P < .05$.

#### Long-term (24 h) retention (Table S9)

Long-term retention was tested only in the subsample that returned at 24 h ($N = 17$), so these correlations are exploratory and uncorrected. No neurophysiological change score was associated with long-term retention in either condition (all $P > 0.16$; Table S9).

**Table S9: Exploratory long-term (24 h) retention correlations.**

| Variable | $\rho$ | 95% CI | $P$ | LOO Range |
|---|---|---|---|---|
| **Panel A. Long-Term Retention With Feedback** ($N = 17$) | | | | |
| *Trained Muscle (APB)* | | | | |
| ΔSDTC | −.154 | [−0.586, 0.387] | .553 | [−0.25, 0.00] |
| ΔRheobase | −.086 | [−0.606, 0.430] | .744 | [−0.21, 0.03] |
| ΔMEP | −.005 | [−0.506, 0.501] | .989 | [−0.10, 0.16] |
| *Untrained Muscle (ADM)* | | | | |
| ΔSDTC | .245 | [−0.312, 0.697] | .342 | [0.11, 0.38] |
| ΔRheobase | −.176 | [−0.670, 0.380] | .497 | [−0.34, −0.07] |
| ΔMEP | −.137 | [−0.629, 0.401] | .599 | [−0.26, 0.00] |
| **Panel B. Long-Term Retention Without Feedback** ($N = 17$) | | | | |
| *Trained Muscle (APB)* | | | | |
| ΔSDTC | −.150 | [−0.611, 0.410] | .566 | [−0.24, 0.00] |
| ΔRheobase | −.230 | [−0.612, 0.290] | .372 | [−0.38, −0.16] |
| ΔMEP | .176 | [−0.337, 0.607] | .497 | [0.09, 0.38] |
| *Untrained Muscle (ADM)* | | | | |
| ΔSDTC | .350 | [−0.201, 0.754] | .168 | [0.24, 0.51] |
| ΔRheobase | −.353 | [−0.718, 0.173] | .165 | [−0.56, −0.29] |
| ΔMEP | −.017 | [−0.525, 0.491] | .951 | [−0.19, 0.11] |

*Note:* Zero-order Spearman correlations ($\rho$) between neurophysiological change scores and 24 h retention with (Panel A) and without (Panel B) visual performance feedback; both muscles, $N = 17$. ΔMEP is the change in MEP amplitude averaged across pulse widths (30, 60, 90 and 120 $\mu$s). Given the exploratory nature and limited power, $P$-values are uncorrected; no association reached $P < .05$. 95% CIs from 10,000 bootstrap resamples; LOO range, leave-one-out coefficient range.
